# Rhinovirus as a Driver of Airway T-Cell Dynamics in Children with Severe Asthma

**DOI:** 10.1101/2024.11.15.623877

**Authors:** Naomi Bryant, Lyndsey M. Muehling, Kristin Wavell, W. Gerald Teague, Judith A. Woodfolk

## Abstract

Severe asthma in children is notoriously difficult to treat, and its immunopathogenesis is complex. In particular, the contribution of T cells and relationships to anti-viral immunity, remain enigmatic. Here, we coupled deep phenotyping with machine learning methods to resolve the dynamics of T cells in the diseased lower airways, and examined rhinovirus (RV) as a driver. Our strategy revealed a T-cell landscape dominated by type 1 and type 17 CD8+ signatures. Interrogation of phenotypic relationships coupled with trajectory mapping identified T-cell migratory and differentiation pathways spanning the blood and airways that culminated in tissue residency, and included transitions between type 1 and type 17 tissue-resident types. These T-cell dynamics were reflected in cytokine polyfunctionality *in situ*. Use of machine learning to cross-compare T-cell populations that were enriched in the airways of RV-positive children with those induced in the blood after RV challenge in an experimental infection model, precisely pinpointed RV-responsive signatures that mapped to T-cell differentiation pathways. Despite their rarity, these signatures were detected in the airways of uninfected children. Together, our results underscore the aberrant nature of type 1 immunity in the airways of children with severe asthma, and implicate an important viral trigger as a driver.

## Introduction

Asthma affects ∼7% of children in the United States, amounting to ∼4.6 million cases (1). Its characteristic recurrent episodes of cough, wheeze, and shortness of breath reflect underlying chronic inflammatory processes in the lower airways and lung parenchyma (2,3). While many children who undergo treatment attain symptom control with minimal exacerbations, ∼5%-10% of cases are severe and remain refractory to standard therapies, including inhaled corticosteroids (3). Consequently, patients with severe disease experience frequent acute exacerbations, often requiring hospitalization, and thus, account for a disproportionate amount of the enormous health and economic burden of asthma (4–6). Despite the advent of biologic therapies that target type 2 inflammatory pathways, and their approval for use in pediatric asthma, their efficacy in children who lack type 2 biomarkers is limited (7,8). Thus, there is a huge unmet need to resolve the immunological basis of severe disease in order to inform new treatments.

T cells are critical to asthma pathogenesis, as exemplified by extensive data to support their contributions to type 2 inflammatory pathways of allergic asthma (9–13). However, recent work by our group and others implicates type 1 responses orchestrated by IFN-γ in promoting the pathogenesis of severe asthma, thereby deviating from the paradigm of “type 2” disease (14–19). Specifically, analysis of cells and mediators in bronchoalveolar lavage (BAL) fluid from children and adults with severe asthma revealed a type 1 bias, including links between type 1- like T cells expressing the chemokine receptor CCR5, and worse lung function (14,19).

Respiratory viruses are likely to be instrumental in this regard, given that they are the most important trigger of wheeze exacerbations in children, and their ability to amplify type 1 responses, including CCR5+ virus-specific T cells, in asthma (20–25). Notably, rhinovirus (RV) accounts for up to 60% of acute episodes in severe asthma, and its detection in ∼ 30% of asymptomatic children with severe disease, implicates this important pathogen as a driver of airway inflammation (5,22,26–28).

Despite these observations, little is known about the nature of T cells that populate the lower airways of children with severe asthma, or how viral infections in early life shape this landscape. Moreover, whether corticosteroid treatment reinforces the type 1 response remains an ongoing debate (29). Adding to the complex picture, pathogenic processes in the pediatric airways likely occur on a backdrop of type 1 immune surveillance inherent to the mucosal barrier, as well as maturation of the immune system. These facets along with the logistics of sampling the diseased airways in children, and challenges to pinpointing rare virus-specific T cells in the lungs, have made it difficult to resolve pathogenic T-cell features in severe asthma.

Here, we aimed to unravel these facets in order to define T-cell based mechanisms governing type 1 responses in the airways of children with severe asthma. This was done by leveraging matched BAL and blood specimens from a highly characterized cohort of children with treatment-refractory recurrent wheeze, and employing novel deep phenotyping and machine learning pipelines. In so doing, we were able to resolve complex type 1-related T cell dynamics with exquisite detail, their relationships to RV. Additionally, we revealed discrete T-cell migratory pathways spanning the blood and airways that culminated in tissue residency, as well as pathways indicating transitions between discrete T-cell functional populations *in situ* that contain RV-responsive T cells. Together, our findings spotlight the aberrant nature of type 1 immunity in the airways of children with severe asthma, and the integral role of RV as a driver.

## Results

### Characteristics of study participants

Study participants included 32, predominantly male children, ages 1-17 years with treatment-refractory recurrent wheeze (**Table 1**). All had failed standard therapy and thus underwent clinically indicated diagnostic bronchoscopy. Children with symptoms of an acute respiratory infection were re-scheduled. Criteria for treatment failure included persistent troublesome symptoms, recurrent unscheduled health care access for wheeze, and/or persistent airflow limitation. At the time of bronchoscopy, 25 patients (78%) were on controller medications, including 24 (75%) who received daily inhaled corticosteroids (ICS). Use of non-steroidal controllers increased with age (p=0.02). Four patients received type 2 biologics (Dupilumab (anti-IL-4Rα), Mepolizumab (anti-IL-5), or Omalizumab (anti-IgE)), and 3 were on oral prednisone. Most patients had evidence of a past or recent infection, based on a history of pneumonia (67.7%) or detection of respiratory pathogens in the BAL fluid (53.1%), respectively. Among those with respiratory pathogens, 13 patients tested PCR-positive for viruses, and RV was most commonly detected (n=10). Bacteria were detected in 10 patients, with cultures identifying *Haemophilus influenza, Moraxella catarrhalis, Pseudomonas aeruginosa, Staphylococcus aureus,* and *Streptococcus pyogenes*. Thus, despite being asymptomatic for infection, detection of respiratory pathogens, and RV in particular, was prevalent in children with recurrent wheeze.

**Table 1.**
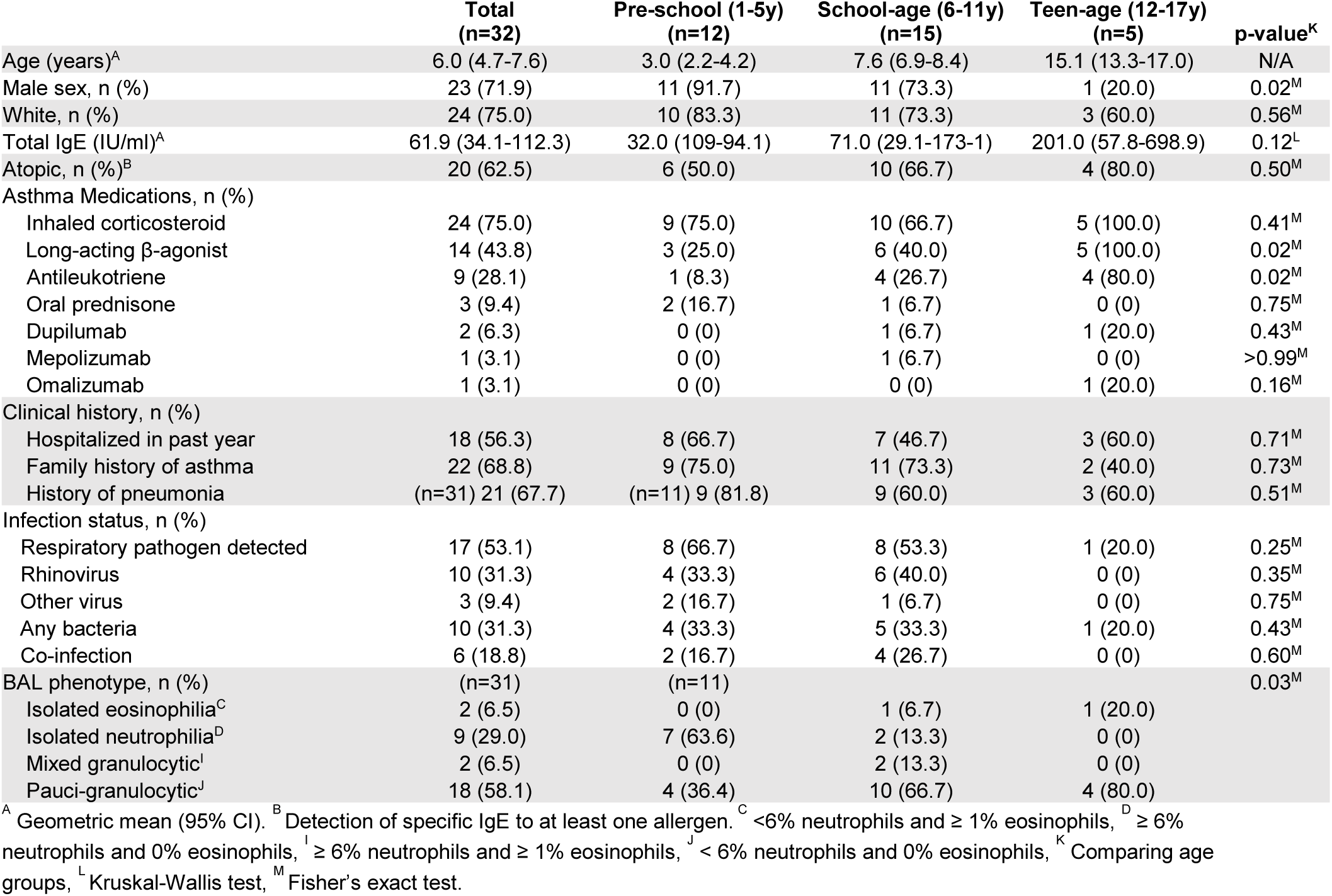
Pediatric patient characteristics.

### Mixed type 1 and type 17 signatures dominate the T-cell landscape in the diseased airways

We first sought evidence for recruitment and antigen priming of, type 1-like T cells in the airways of children with treatment-refractory wheeze, by comparing matched BAL and blood specimens. As expected, spectral flow cytometry analysis (**Supplemental Table 1**) revealed enrichment of CD8+ T cells in the lungs (30,31), and this was most marked in patients with one or more respiratory pathogen (mean CD4:CD8 0.5 ± 0.07 vs. 0.8 ± SEM 0.1, p=0.03; **Figure 1A, B and C**). The majority of T cells in the airways displayed a memory phenotype (>90%). Both CD4+ and CD8+ effector memory T cells (T_EM_, CD45RO+CCR7-) were markedly enriched in the BAL versus the blood (72.3% ± 2.3 vs. 7.7% ± 0.7 and 86.1% ± 1.8 vs. 17.4% ± 1.4, respectively), whereas frequencies of more differentiated TEMRA-like cells (CD45RO-CCR7- CD27-) were low and similar at both tissue sites (**Figure 1D and Supplemental Figure 1A**). As we reported previously (19), most CD4+ and CD8+ T cells in the airways expressed CCR5 (86.0% ± 1.4 and 83.7% ± 2.1, respectively), which is notable for its role in memory responses to respiratory viruses (**Figure 1E**) (32,33). Probing classical markers of type 1 (CCR5, CXCR3), type 2 (CCR4, CRTH2), and type 17 (CCR4, CCR6, CD161) T-cell types indicated the accumulation of type 1 and 17 T cells in the airways (**Figure 1E**). While CXCR3 was the most abundantly expressed marker on both memory CD4+ and CD8+ T cells, CCR6 and CD161 were expressed at higher frequencies on CD4+ T cells compared with CD8+ cells. Further resolution of airway memory CD4+ T-cell signatures identified a predominance of type 17 canonical and non-canonical subsets including Th17 cells (CCR6+CD161+CCR4+), “plastic” Th17 or Th17.1 cells (CXCR3+CCR6+CD161+), and Th17-like cells that lacked CCR4 (CCR6+CD161+CCR4-) (**Figure 1F and Supplemental Figure 1B**). Classical Th1 cells (CXCR3+) were less frequent, whereas classical Th2 cells (CCR4+ only) and pathogenic Th2 effectors (Th2a, CRTH2+) were the least abundant. By contrast, Th2 cells were the dominant subtype in blood (**Supplemental Figure 2A**). CD8+ T cell signatures were predominantly type 1 (CXCR3+), with type 17 (CCR6+CD161+) and type 17.1 (CXCR3+CCR6+CD161+) cells also present, albeit at lower frequencies (**Figure 1F and Supplemental Figure 1C**). As expected, the majority of T cells in the airways were tissue-resident (68.5% ± 2.6 and 75.9% ± 2.9 for CD4+ and CD8+ T cells respectively) based on expression of CD69 with or without CD103 (**Figure 1G**), and tissue-resident T cells (T_RM_) expressed higher levels of chemokine receptors as compared with their non-resident counterparts (**Figure 1H**). No differences in the frequencies of memory T-cell types were identified when comparing children with (n=17) or without (n=15) a respiratory pathogen, or on the basis of ICS use (**Supplemental Figures 2, B and C**) (34–36). Although cell frequencies were similar regardless of atopic status (**Supplemental Figure 2D**), levels of total IgE among atopic patients were positively correlated with frequencies of Th1 cells (CXCR3+, r=0.56, p=0.01), and these were inversely correlated with Th17-like cells (**Supplemental Figure 3**). Analysis of T cells by age implied preferential accumulation of type 1-like CD4+ and CD8+ cells in the airways versus blood (**Supplemental Figure 4**). Among the subset of patients for whom lung function tests were performed (n=16), no relationships were identified with T-cell types (data not shown). Together, these results supported the priming and enrichment of type 1 and type 17 signatures in the diseased airways regardless of ICS treatment, and a paradoxical relationship to IgE.

**Figure 1.**
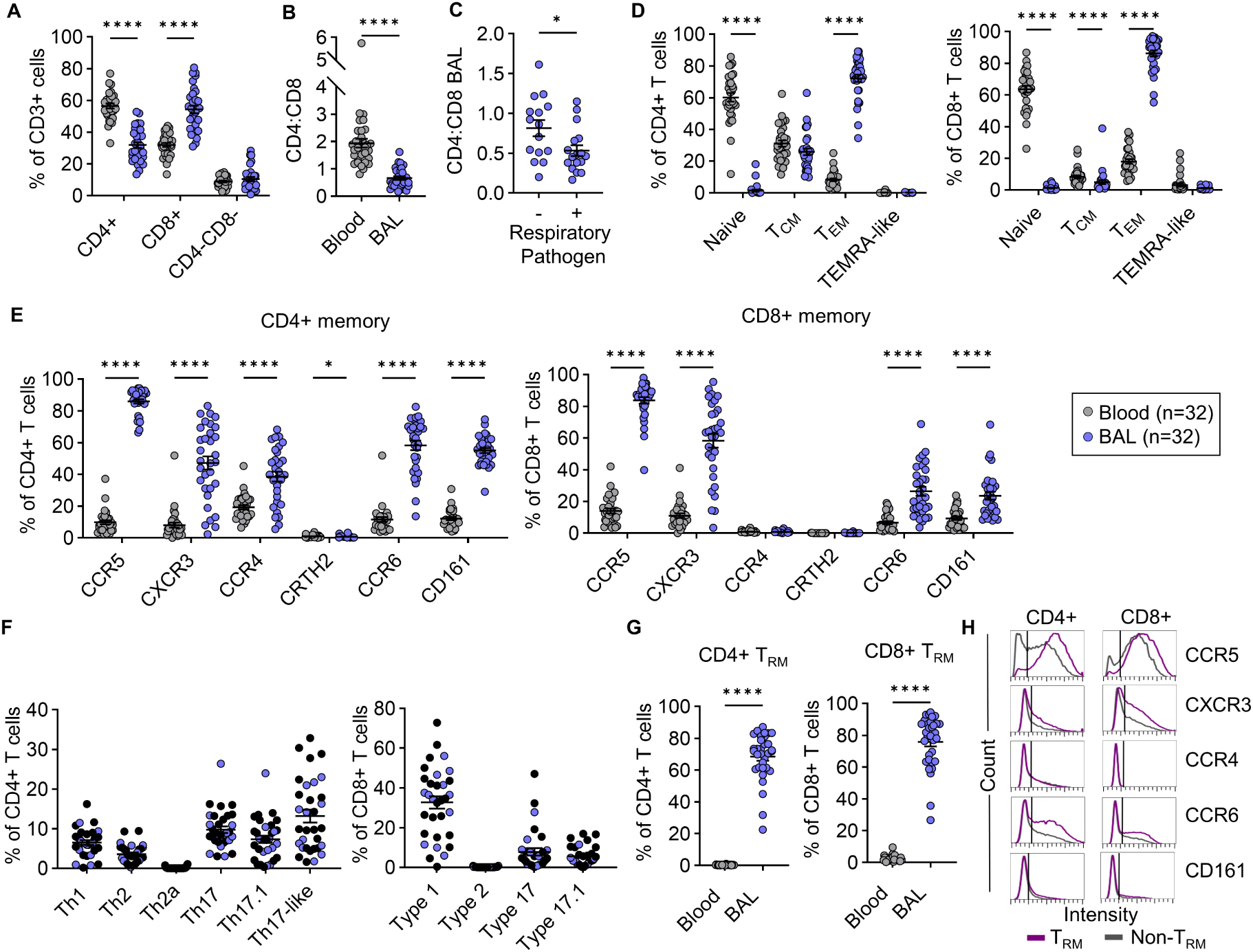
Mixture of type 1 and 17 signatures dominate the T-cell landscape in the lower airways of children with recurrent wheeze. (A) T cell frequencies in blood and BAL, as a percentage of CD3+ cells. (B) CD4:CD8 ratio in blood and BAL. (C) BAL CD4:CD8 ratio in patients with (n=17) or without (n=15) a respiratory pathogen. (D) Naïve (CD45RO-CCR7-), central memory (CD45RO+CCR7+, T_CM_), effector memory (CD45RO+CCR7-, T_EM_), and terminal effector memory-like (CD45RO-CCR7-CD27-, TEMRA-like) T cells in blood and BAL, as a percentage of CD4+ or CD8+ T cells. (E and F) Phenotype of memory (T_CM_, T_EM_, and TEMRA-like) CD4+ or CD8+ T cells in blood or BAL. Black symbols denote atopic patients (n=20) in F. (G) Frequency of CD4+ or CD8+ tissue-resident memory T cells in blood and BAL (CD69+CD103- and CD69+CD103+, T_RM_). (H) Histograms depicting the median expression of surface receptors on CD4+ and CD8+ Trm and Non-T_RM_ in BAL. Horizontal bars denote mean ± SEM. Multiple Wilcoxon tests with Holm-Sidak correction (A, D, & E) and Wilcoxon matched-pairs signed rank test (B, C, & G) calculated statistical significance. *p<0.05, **p≤0.01, ***p≤0.001, ****p≤0.0001.

### Aberrant T-cell dynamics in the airways link to infection status

To better understand the complexity and dynamics of T cells in the diseased airways, we next leveraged the high-dimensional nature of our data. Data visualization using density maps confirmed markedly different T-cell distributions between airways and blood, owing primarily to expression of CCR5 and T_RM_ markers (CD69 and CD103) on most airway cells (**Figure 2, A and B**). Clustering analysis by PhenoGraph yielded 68 distinct cell clusters across BAL and blood (**Figure 2C**) (37). Hierarchal clustering delineated two sets of cell clusters that were enriched in the airways versus the blood (groups #1 and #5) and comprised mostly CD4+ and CD8+ T_EM_ (CD45RO+CCR7-) (**Figure 2D**). Most CD4+ T_EM_ clusters in group #1 were T_RM_ CXCR3^lo^, and expressed the activation markers PD-1, ICOS, and CD95 (**Figure 2D and Supplemental Figure 5**). Expression of the type 1-orchestrating transcription factor T-bet was low, consistent with its downregulation in tissues (38,39). Active responses were evidenced by the presence of robustly proliferating (CD38+Ki-67+) cells (cluster #33 & 58). In addition to Th1- like clusters (CXCR3+T-bet+), memory CD4+ T_EM_ unique to the airways included CD69+ types that were CD161+ or CCR6+CD161+ (#3, 5, 28, 42, 48, 58). Low co-expression of Th1 markers by some of these subtypes indicated Th17.1 phenotypes, suggesting transitions between type 1 and type 17 functional subsets (40,41) (**Figure 2D and Supplemental Figure 5**). Additional non-resident CD4+ T_EM_ phenotypes were shared with those found in the blood, including Th17- like cells (CD69-CD103-CCR6+CD161+: clusters #31 & 32) that displayed increased stemness (TCF-1+) and were less activated (CD127+), perhaps reflecting recent egress from blood (42,43). Evidence of T-cell transitions was supported by using PHATE analysis to infer a T-cell trajectory that linked cluster #32 with type 1 (#39) and the type 17-related (#3, 5, 28, and 42) T_RM_ clusters (**Figure 3, A, B, and C**) (44). Specifically, transitions along the path from cluster #32 (enriched in the blood) to cluster #3 (exclusive to the BAL) were associated with decreased expression of the differentiation markers CD27 and TCF-1, and a concomitant increase in tissue-resident and activation markers (CD69, CD103, and PD-1) (45). Together, these features indicated T-cell migration from the blood, establishment of tissue residency, and functional transitions in the tissue.

**Figure 2.**
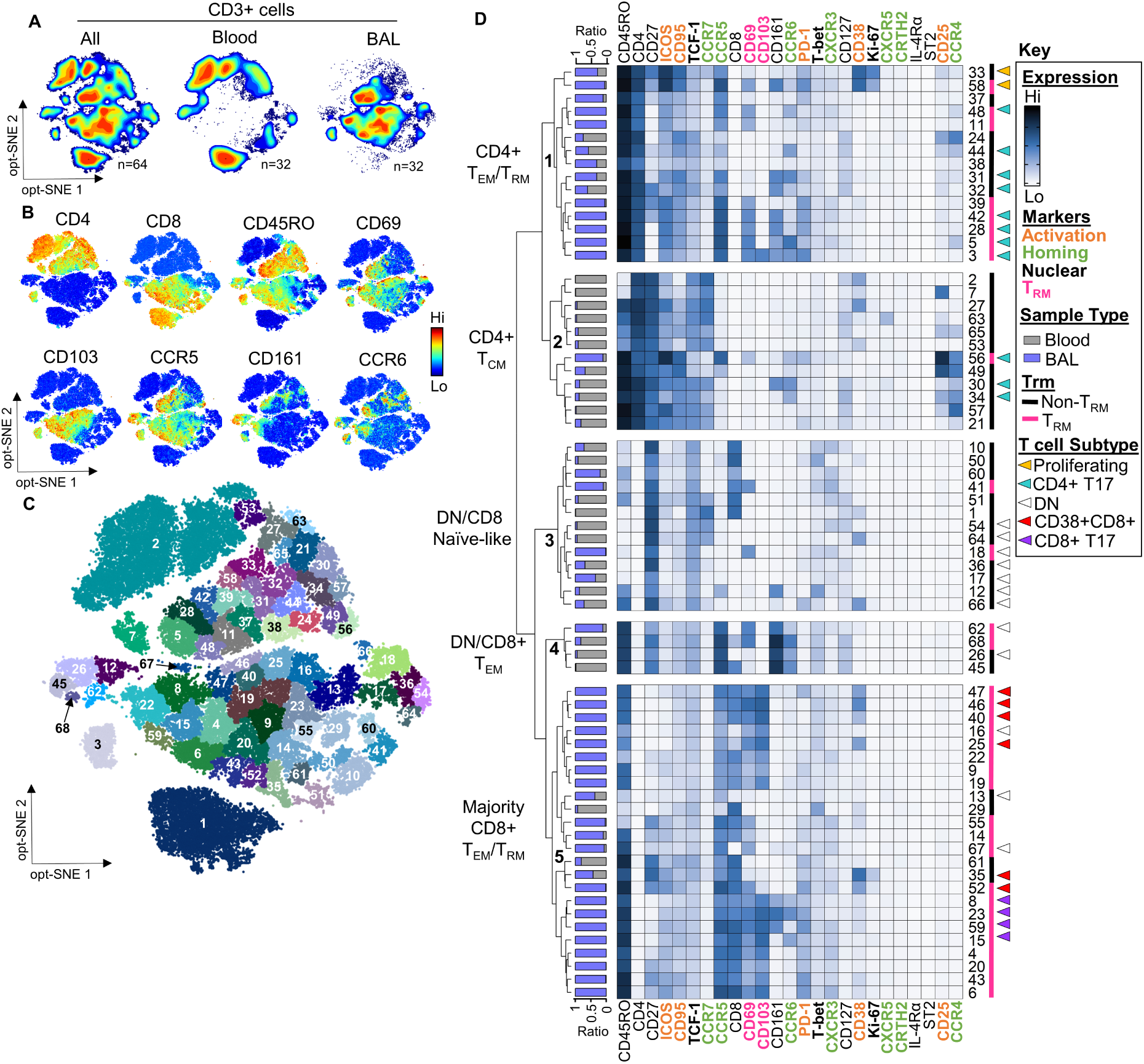
The T cell landscape in the lower airways of children with recurrent wheeze is highly dynamic. (A) Density maps showing the distribution of blood and BAL T cells on opt-SNE axes. (B) Expression of select phenotypic markers on opt-SNE axes. (C) PhenoGraph clusters projected on opt-SNE axes. (D) Heatmap showing the median expression of each marker, for clusters generated by PhenoGraph. Annotation on the left denotes the proportion of cells with each cluster that come from blood or BAL

**Figure 3.**
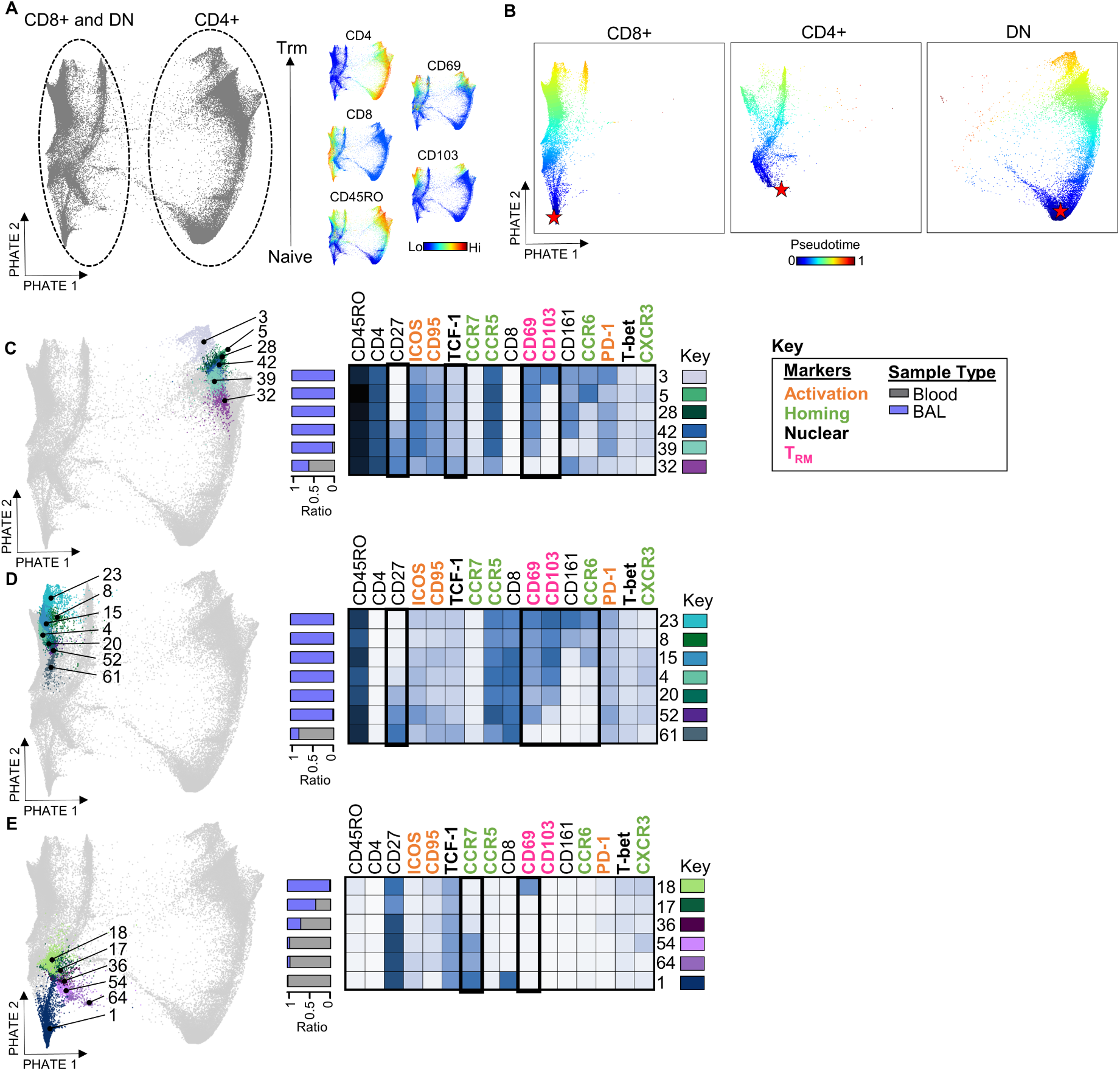
PHATE analysis captures transitions in T cell clusters in the lower airways. (A) PHATE map generated on CD3+ cells from matched blood and BAL (n=32). Heatmaps showing the distribution of select markers on PHATE axes. (B) Pseudotime trajectory of CD4+, CD8+, and DN T cells projected on PHATE map. The star denotes the starting point for wishbone analysis. Select PhenoGraph clusters projected on PHATE map to show transitions within (C) CD4+, (D) CD8+, and (E) DN T cells. Black boxes indicate markers that are differentially expressed between clusters. Left-hand annotation denotes the proportion of cells within each cluster that come from blood or BAL (D-F).

CD8+ T_RM_ dominating the T-cell landscape in the airways displayed a predominant type 1 signature (CXCR3+T-bet+) (**Figure 2D**). Expression of CCR6 or CD161 was limited to four T_RM_ clusters (#8, 15, 23, 59) that co-expressed variable type 1 markers. PHATE analysis mapped these type 17-related T_RM_ to the same pathway as type 1 T_RM_ (clusters #4, 20, 52) and non-T_RM_ (cluster #61) signatures, further supporting transitions between type 1 and 17 CD8+ T cells in the lower airways (**Figure 3D**). Again, expression of PD-1, ICOS, and CD95 was a feature within CD8+ T_RM_, although levels and combinations varied across cell clusters (**Figure 2D and Supplemental Figure 5**). High expression of the activation marker CD38 on six CD8+ T-cell clusters was notable for its link to chronic viral infection (46). These included a subset of proliferating non-resident type 1 cells shared with blood (cluster #35) and its CD69+ counterpart unique to the lungs (#52), as well as T_RM_ types that lacked ICOS and PD-1 (#25, 40, and 46).

Interspersed among CD8+ T_EM_ on the heatmap were three double-negative (DN, CD4-CD8-) clusters that were enriched in, or unique to, the lungs (#13, 16 & 67). Also, unique to the lungs was a fourth CD69+ DN subset (#18) that resembled naïve “effector-like” cells for its Th1 differentiation (CXCR3) despite a naïve phenotype (CD45RO-CCR7-CD27+TCF-1+) (47). Trajectory mapping showed cluster #18 at the intersection of naïve DN T cells (clusters #54 and 64) and naïve CD8+ T cells (cluster #1), pointing to convergent pathways (**Figure 3E**).

High expression of activation markers on T_RM_ cells in the airway lumen indicated the presence of activated T cells in inflamed lung tissue niches. Expert gating on memory CD4+ and CD8+ T cells that were PD-1+ICOS+CD95+ revealed higher frequencies in patients with respiratory pathogens versus those without (57.6% ± 2.3 vs 46.6% ± 2.7, p=0.009 for CD4+ T cells; and 13.1% ± 1.9 vs 8.6% ± 1.6, p=0.09 for CD8+ T cells). Importantly, this relationship was maintained after adjusting for changes in frequencies of activated T cells with age (p=0.05 for CD4+ T cells; and p=0.04 for CD8+ T cells) (**Figure 4, Supplemental Figure 6**). Together, these results demonstrated aberrant T-cell dynamics in the diseased airways and their link to respiratory infections.

**Figure 4.**
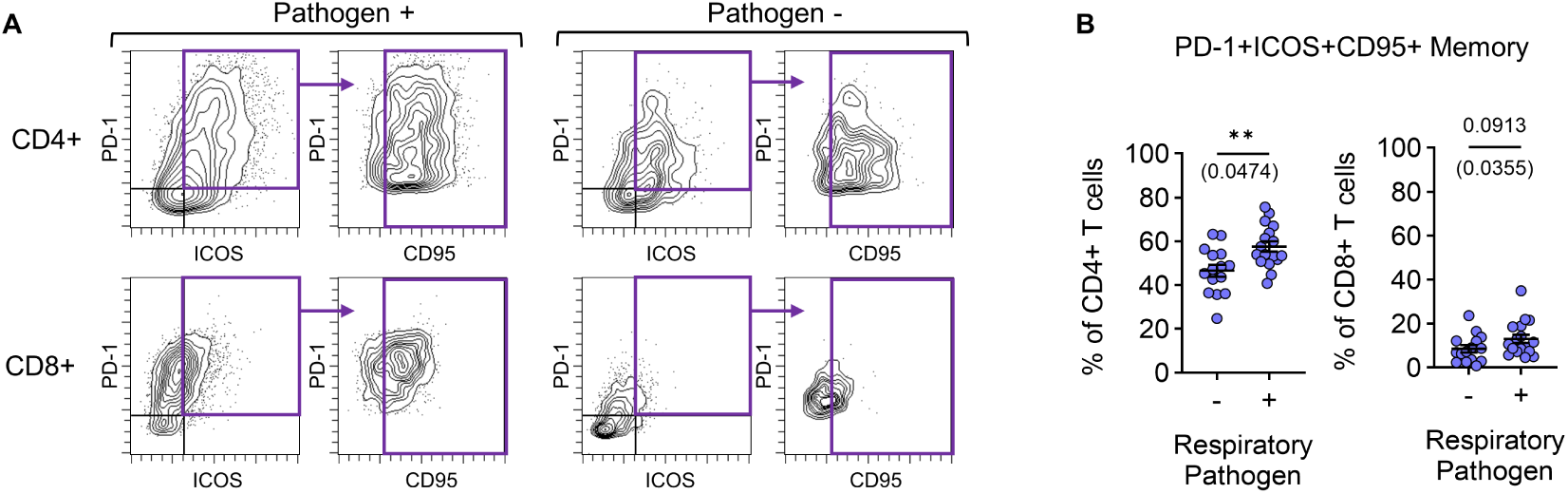
Respiratory pathogens are linked to increased T cell activation in the lower airways. (A) Representative scatter plots showing gating of PD-1+ICOS+CD95+ T cells within total memory (T_CM_, T_EM_, TEMRA-like) T cells. (B) Comparison of PD-1+ICOS+CD95+ memory T cells in the BAL, as a frequency of CD4+ or CD8+ T cells in children with (n=17) and without (n=15) a respiratory pathogen. P values in parentheses are corrected for age. Horizontal bars denote mean ± SEM. Mann-Whitney test and multiple logistic regression used to calculate significance (B). **p≤0.01

## Rhinovirus-related signatures contribute to aberrant T-cell dynamics

We next asked whether RV, a major instigator and exacerbator of childhood wheeze, might act as a driver of T-cell changes in the lungs. The computational tool T-REX was used to pinpoint candidate virus-specific signatures, which we hypothesized would be enriched in the airways versus blood, but nonetheless present at very low frequencies (48). T-REX is ideally suited to detect rare cell types based on the degree of change in discrete complex signatures between samples. T cells were compared between patient groups with and without RV infection based on PCR positivity (RV+: n=10, and RV-negative: n=22) (**Supplemental Table 2**). While demographic and clinical characteristics were similar between groups, frequencies of airway neutrophils were increased in the RV+ group (median, 29.0% vs 2.5%; p=0.03), and bacterial co-infection was more common (40.0% vs 9.1%, p=0.06), similar to what has been reported previously (49,50). Three subjects in the RV-negative group tested positive for adenovirus (n=1) or metapneumovirus (n=2), whereas RV+ patients tested negative for other viruses. T-REX analysis of airway T cells, coupled with marker enrichment modeling (MEM) to assign complex signature labels, identified four dominant signatures that were enriched in the RV+ group versus the RV-negative group (**Figure 5, A and B**). By contrast, nominal differences were observed in the blood (**Supplemental Figure 7A**). T-REX populations 1-3 were memory CCR5+CD4+ T cells that expressed high levels of ICOS and activation markers (CD38, PD-1, and CD95). T-REX 1 mapped to PhenoGraph cluster #33, displayed robust proliferation (Ki-67^hi^), and constituted a resident Th1 phenotype (CD69+CXCR3+T-bet+CD27+) that was CD25+ (**Figure 5A and Supplemental Figure 7, B and C**). T-REX 2, which also mapped to PhenoGraph cluster #33, was a non-resident (CD69-) counterpart of T-REX 1 displaying lower proliferation, consistent with a precursor that had not yet acquired tissue residency. T-REX 3 was a resident Th17-like population (CD69+CD161+CCR6^lo^) that mapped to PhenoGraph cluster #58, whereas T-REX 4 contained CD8+ T_RM_ (CD38+PD-1+) mapping to cluster #47. Notably, frequencies of T-REX 2 and 3 correlated with neutrophil frequencies in the airways (**Figure 5D and Supplemental Table 2**). Projection of T-REX 1-3 onto trajectory pathways (**Figure 3A**) further highlighted the relatedness of T-REX 1 and 2, as well as the advanced differentiation (CD27^lo^) of T-REX 3 (**Supplemental Figure 7D**). Overlay of phenograph clusters on the PHATE map also showed “less-activated” (CD38-) T_RM_ counterparts of T-REX 1/2 (cluster #39) and T-REX 3 (clusters #28 & 42), as well as a blood precursor (cluster #32) (**Supplemental Figure 7E**). “Less-activated” (clusters #4 & 20) and precursor (cluster #61) counterparts for the CD8+ population, T-REX 4, were also identified (**Supplemental Figure 7F**). Notably, these counterparts were present at appreciable frequencies in both RV- and RV+ patients (**Supplemental Figure 7G**).

**Figure 5.**
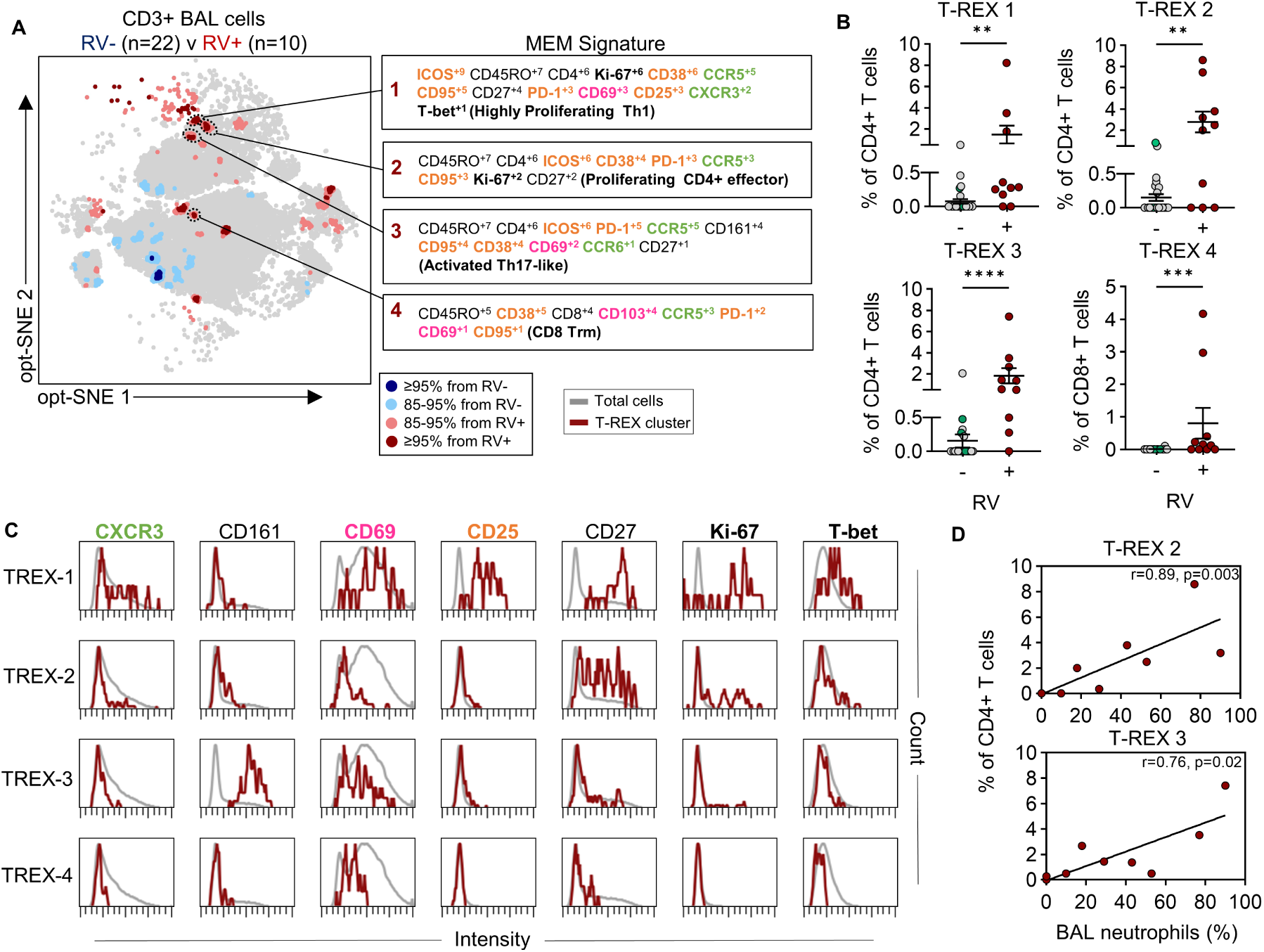
Highly activated CCR5+ T cells are linked to RV infection in the lower airways. (A) T-REX plot comparing BAL T cells from RV-(n=22) and RV+ (n=10) patients. Red populations are up in the RV+ group, and blue are up in the RV-group. Marker enrichment modeling (MEM) was used to characterize dominant populations. Each marker is scored on a scale of 1-10 based on its enrichment in the cluster. (B) Frequency of T-REX populations in RV- and RV+ patients. Green data points are patients positive for other viruses (n=3). (C) Histograms showing the median expression of select markers on T-REX populations. (D) Spearman correlations of T-REX population frequency in RV+ subjects vs. BAL neutrophil frequency (n=9). Horizontal bars denote mean ± SEM. Line denotes linear regression. Mann-Whitney used to calculate statistical significance (B). *p≤0.05, **p≤0.01, ***p≤0.001, ****p≤0.0001.

Five additional “highly variable” populations (T-REX 5-9) were enriched in the airways of RV+ patients, although frequencies varied among RV+ patients, with each population containing >50% of cells from a single patient (**Supplemental Figure 8**). These included a CD161+ memory CD4+ signature shared with blood that mapped to PhenoGraph cluster #32 (T-REX 5), and three resident CD8+ effector populations (T-REX 6-8) that were unique to the lungs, and expressed variable levels of CD38, PD-1, and CD95 (mapping to clusters #19, 43, 47). T-REX 9 corresponded to the only DN cluster that was unique to the airways (#18), and notable for its naïve “effector-like” signature. Further analysis of the frequencies of T-REX populations 1-9 confirmed their marked enrichment in BAL versus blood in the RV+ group, and found no differences based on the presence of bacterial pathogens (**Supplemental Figure 9**). Together, these results pointed to the expansion and persistence of RV-related T-cell signatures in the diseased airways, and their contribution to aberrant T-cell dynamics.

### T-REX populations in the diseased airways display RV-responsive hallmarks

To further probe whether RV-related signatures in the airways were virus-specific, we next analyzed their similarity to RV-specific T cells whose expansion peaks on day 7 in the blood after RV challenge (23). We previously confirmed the ability for T-REX analysis to detect these cells with the same precision as MHCII/peptide tetramer staining (48). Applying T-REX analysis to specimens from RV-challenged adults with and without asthma revealed expansion of multiple CD4+ and CD8+ populations after infection (**Figure 6, A and B**, **Supplemental Figure 10**). Among these, only one CD4+ T_EM_ population was uniformly expanded on day 7 (average 11-fold versus baseline). This population (designated T-REX A) showed strong proliferation (Ki-67^hi^), was armed for lung-homing (CCR5+), highly activated (ICOS^hi^CD38^hi^CD95^hi^CD25^mid^PD-1^mid^), and displayed Th17.1 features (CXCR3+CD161+CCR6^lo^T-bet+) (**Figure 6, C and D**). Using root mean square deviation (RMSD) analysis to quantitatively compare the MEM label of T-REX A to signatures enriched in children’s airways, found high similarity with T-REX 1 (proliferating Th1 T_RM_), T-REX 2 (proliferating CD4+ effector), and T-REX 3 (Th17-like T_RM_) (94.6%, 91.9%, and 90.1% respectively) (**Figure 6E**) (51). These data imply that RV-related signatures in the diseased airways of RV+ children are virus-specific, and poised to undergo viral activation and phenotypic transitions.

**Figure 6.**
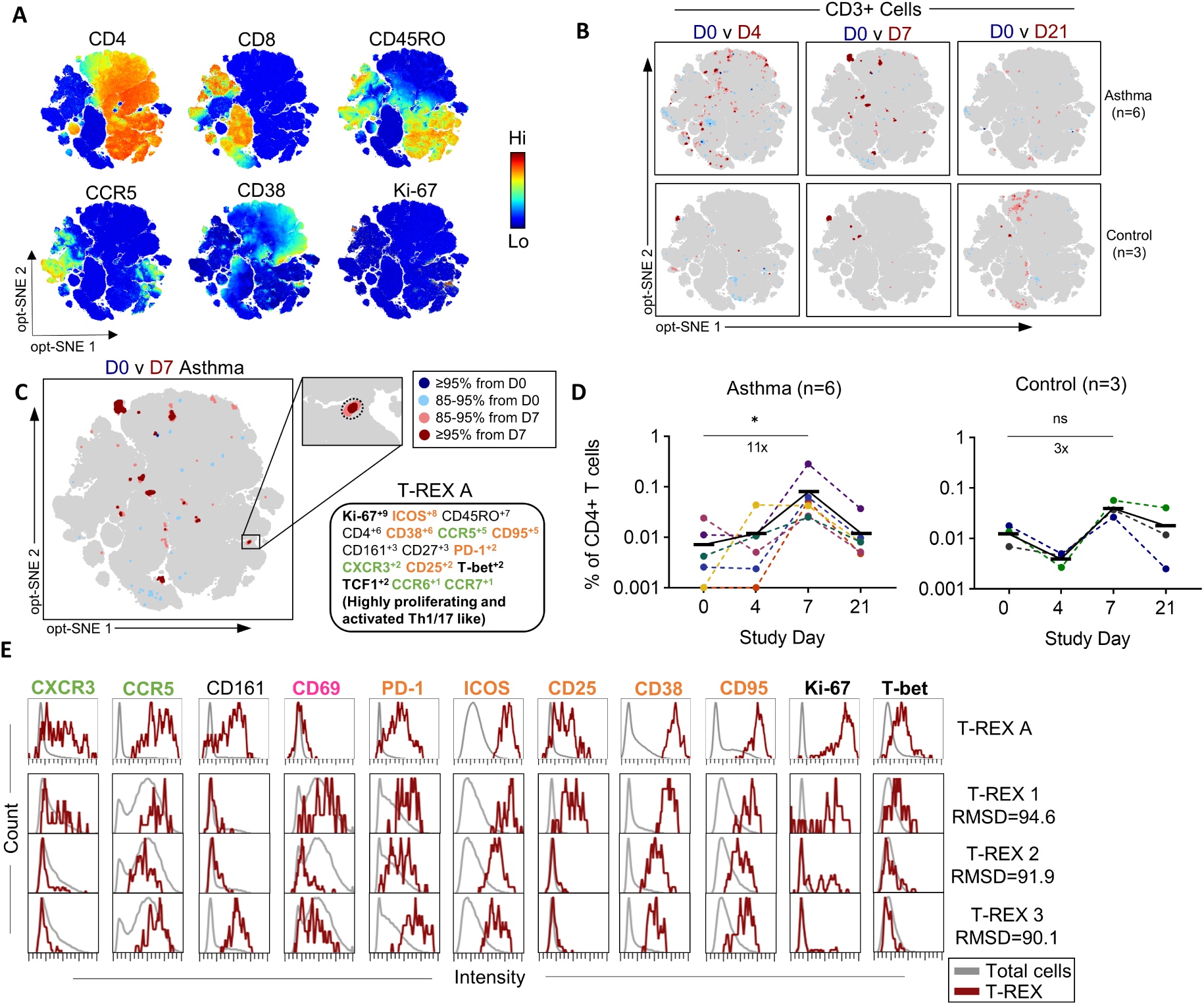
T cell signatures enriched in RV+ children express RV-responsive hallmarks. (A) Heatmaps showing the distribution of marker expression on opt-SNE axes. (B) T-REX’s comparing days 4, 7, and 21 of RV infection to day 0 in adults with asthma and healthy controls. (C) MEM signature of T-REX A. (D) Quantification of T-REX A frequency at each study day. Numbers denote the average fold change between days 0 and 7. Lines denote the mean value for each study day. (E) Histograms comparing the T cell signature identified in adults to those enriched in the BAL of RV+ children. Root mean square deviation (RMSD) shows similarity to of T-REX 1-3 to T-REX A. Friedman test with Dunn’s correction for multiple comparisons calculated statistical significance (D). * p<0.05.

### Multi-functional and cytotoxic CD4+ and CD8+ T cells populate the diseased airways

To test pro-inflammatory effector functions of T cells in the airways, we analyzed an array of intracellular cytokines in BAL T cells from six children with recurrent wheeze (**Supplemental Tables 3 and 4**). Most cytokine-positive CD4+ memory T cells were IFN-γ+ (48.6% ± 5.6) and TNF-α+ (44.3% ± 7.7), and around one third were IL-2+ (30.4% ± 4.6) (**Figure 7A**). Lower frequencies of cytotoxic (GzmB+) CD4+ (7.7% ± 2.5), Th17 (IL-17A+, 7.5% ± 1.2) and Tfh-like cells (IL-21+, 5.4% ± 1.5) were also identified, whereas Th2-like cells (IL-4+, IL-5+, IL-13+) were infrequent. In contrast to CD4+ T cells, memory CD8+ T cells were primarily IFN-γ+ (66.8%±5.1) and GzmB+ (51.9% ± 5.1), whereas frequencies of TNF-α+, IL-2+ and IL- 17A+ cells were lower (23.4% ± 5.6; 27.8% ± 5.9; and 6.0% ± 2.1 respectively) (**Figure 7A**).

**Figure 7.**
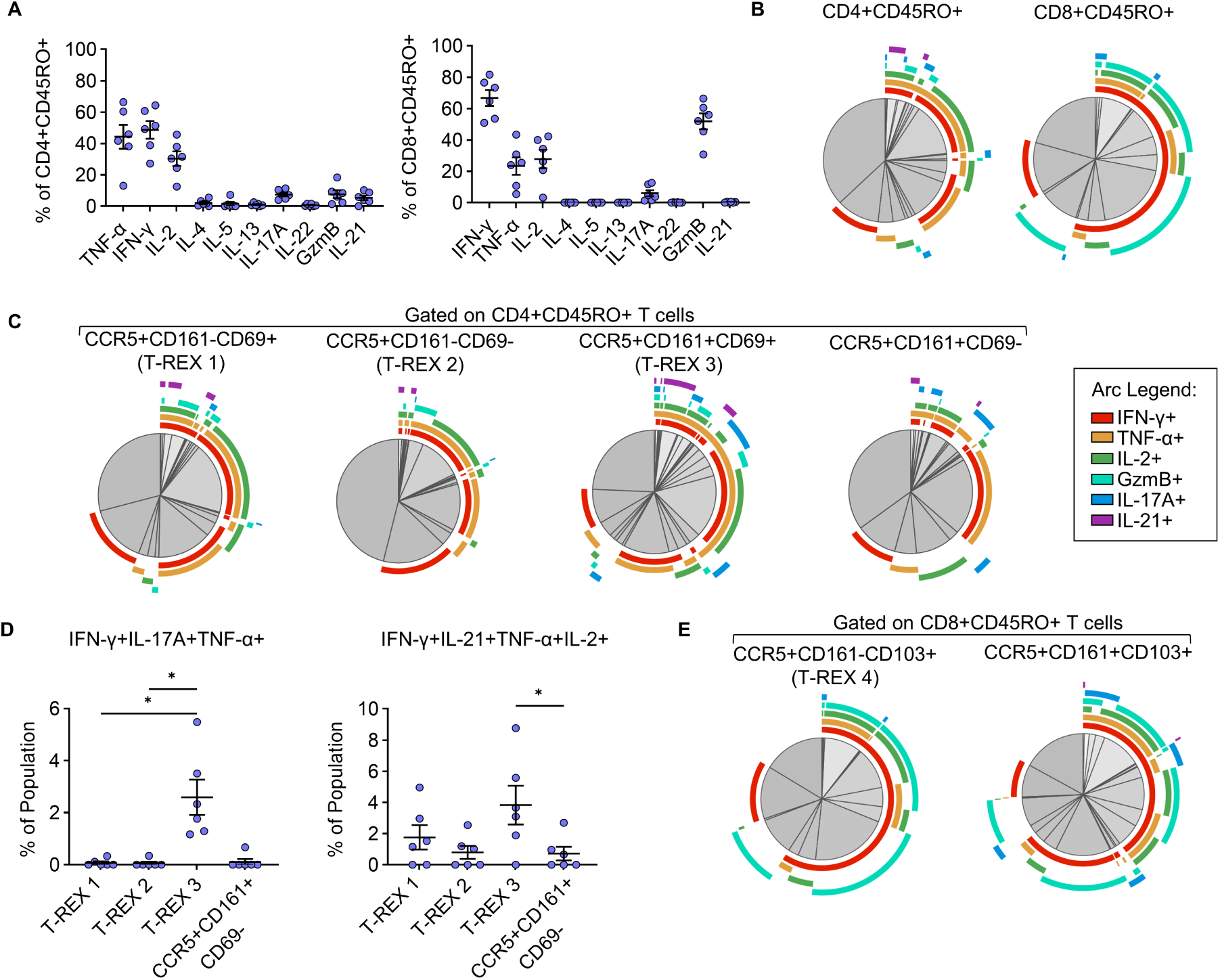
Polyfunctional T cells populate the lower airways of children with refractory wheeze. (A) Cytokine positive cells in the airways as a frequency of CD4+ and CD8+ memory T cells (n=6). (B) Spice plots showing averages of cytokine combinations within CD4+ and CD8+ memory T cells. (C) Spice plots showing averages of cytokine combinations within CD4+ T-REX related populations. Non-T_RM_ (CD69-) counterpart of T-REX 3 is shown as comparison. (D) Frequency of select SPICE plot populations within each T-REX related population shown in panel C. (E) Spice plots showing averages of cytokine combinations within a CD8+ T-REX related population. CD161+ counterpart of T-REX 4 is shown as a comparison. Horizontal bars denote mean ± SEM. Friedman test used to calculate significance (D). * p<0.05.

SPICE analysis (52) of cytokine combinations within CD4+ and CD8+ memory T cells confirmed co-expression of IFN-γ and TNF-α in both CD4+ and CD8+ T cells, as well as triple- and quadruple-positive subsets expressing a combination of IL-2, IL-17A, IL-21, and GzmB (**Figure 7B and Supplemental Figure 11A**).

To analyze effector functions of virus-responsive signatures, we used a MEM-directed gating strategy based on a minimal marker set to identify populations in the cytokine assay that aligned with T-REX populations 1-4 in RV+ children (T-REX 1: CCR5+CD161-CD69+, T-REX 2: CCR5+CD161-CD69-, T-REX 3: CCR5+CD161+CD69+, T-REX 4: CCR5+CD161+CD103+).

The 4-hour stimulation period minimized changes in expression of CCR5, CD69, and CD103; however, CXCR3 was excluded from analyses owing to its downregulation upon stimulation (**Supplemental Figure 11B**). Cytokine profiles of all candidate T-REX populations were dominated by IFN-γ and TNF-α. However, the CD4+ signature resembling T-REX 3 (CCR5+CD161+CD69+) displayed increased polyfunctionality (IFN-γ+TNF-α+IL-17A+ and IFN-γ+TNF-α+IL-2+IL-21+) compared with its CD161- and non-T_RM_ (CD69-) counterparts, indicating enrichment of Th17.1 and T peripheral helper-like (Tph-like) cells within this population (**Figure 7C and D**, **Supplemental Figure 11C**) (53). As expected, the majority of the CD8+ subset resembling T-REX 4 (CCR5+CD161-CD103+) were GzmB+IFN-γ+, and co-expressed TNF-α or IL-2 but had lower IL-17 expression compared with their CD161+ counterparts **(Figure 7E and Supplemental Figure 11D)**. These results confirmed that virus-responsive signatures in the diseased airways are equipped to exert polyfunctional and cytotoxic functions that promote type 1 and type 17 responses.

## Discussion

Although recent studies implicate type 1 responses in the pathogenesis of severe asthma in adults (14,18,54), the T-cell landscape in the airways of children with severe recurrent wheeze remains largely unexplored. Our study used novel deep phenotyping methods and machine learning tools to address this major knowledge gap. Further, we sought to establish a mechanistic link between a major viral trigger and T-cell dynamics in the diseased airways.

Rigorous investigation of T-cell attributes in the context of clinical and demographic variables, demonstrated aberrant T-cell dynamics in the diseased airways that were dominated by complex mixtures of type 1 and type 17 CD4+ and CD8+ T-cell effectors. Integral to these dynamics were migratory pathways spanning the blood and airways that culminated in tissue residency, as well as phenotypic relationships indicating active transitions between type 1 and type 17 functional subsets. By using powerful machine learning methods to detect rare T-cell populations, we defined RV-responsive elements that mapped to T-cell trajectories. To our knowledge, our study is the first to use this approach to establish RV as a potential driver of aberrant T-cell behavior in the airways. Our innovative strategy surmounted barriers to identifying virus-specific T cells whose very low numbers in the airways and unknown epitope specificities preclude identification by labeling with MHCII/peptide tetramers. Based on our collective findings, we propose that a high incidence of viral infections - RV in particular - contributes to aberrant T-cell dynamics, a vicious cycle of T-cell-based inflammation, and corticosteroid resistance, in susceptible children. By helping to solve the conundrum of RV as an instigator and/or driver of disease, our study informs both the immunopathology of severe asthma and potential therapeutic targets to alter the clinical course.

Our results showed that starting from ∼18 months of age, the vast majority of both CD4+ and CD8+ T cells in the airways are CCR5+ T_RM_, indicating early accumulation of memory T cells in lung tissue (55–57). While type 1-related T cells featured prominently, type 2 signatures, a hallmark of allergic asthma, were infrequent. This was borne out by expression of IFN-γ in the majority of cytokine-expressing CD4+ and CD8+ T cells. Notably, a large proportion of airway CD4+ T cells also displayed type 17-related features, as evidenced by their co-expression of CD161 and CCR6 with or without CXCR3 and CCR4, indicating the presence of Th17, Th17.1 and Th17-like varieties. Moreover, expression of IL-17A by both CD4+ and CD8+ T-cell subsets expressing CD161 indicated contributions of bona fide type 17 cells capable of secreting IL-17. Previous studies in children and adults have linked IL-17A in the serum and airways to asthma severity (58–61). In addition, CCR5+CD161+CD4+ T cells have been linked to decreased lung function in adult patients (14). IL-17A exerts its effects on multiple cell types, including airway smooth muscle cells, endothelial cells, and fibroblasts, and is arguably best known in asthma for mediating the recruitment of neutrophils to the lungs (62–67). The interplay between type 17 inflammation and corticosteroids, as well as insensitivity of Th17 cells to corticosteroid-mediated cell death, may serve to compound type 17-mediated lung inflammation in severe asthma (36,68–70).

In addition to CD4+ and CD8+ T-cell types, a novel finding in our study was the presence of DN T cells (CD4-CD8-) in the airways, including a tissue-resident naïve “effector-like” type expressing CXCR3 that was unique to this site. DN T cells in the lungs, which may include γδ T cells (reported to be ∼13% of total CD3+ cells in the lungs of children), respond rapidly to respiratory viruses, and can exert regulatory and effector functions (71,72). As such, these cells warrant further investigation, including their ties to the production of autoantibodies, which have been linked to asthma exacerbations and severe disease (73–76).

Our results revealed a highly dynamic T-cell landscape in the diseased airways as evidenced by a broad spectrum of signatures of activation, proliferation, and tissue residency across multiple related populations. In particular, co-expression of PD-1, ICOS, and CD95, pointed to a high degree of T-cell activation. This was observed on RV-responsive T-cell signatures undergoing robust proliferation, as well as on broader T-cell populations displaying complex type 1 and type 17 signatures. In particular, the presence of Th17.1 cells (CD161+CXCR3+CCR6+) indicated active transitions between type 1 and type 17 cells, a finding bolstered by trajectory mapping, and detection of T-cell populations co-expressing IL-17 and IFN-γ. These data, coupled with migratory pathways spanning blood and tissue, supported a model wherein egress of T cells from the blood culminates in establishment of tissue residency, activation, and differentiation along a type 1/type 17 axis. Pinpointing Th17.1 cells in our study was important since these constitute a polyfunctional and pathogenic subset involved in diverse chronic inflammatory disorders that is also resistant to suppression by regulatory T cells in the inflamed patient (40,77,78). In extension of this notion, the inflamed milieu may also serve to override pathways acting to regulate the functions of lung-resident T cells through PD- 1, ICOS, and CD95, thereby further amplifying the inflammatory cascade (79–82).

Our study does not exclude a role for T-cell exhaustion in perpetuating airway inflammation. This could arise through chronic antigen stimulation and consequent dysregulated responses. Arguing against this, our work identified only low numbers of TEMRA-like cells (CD45RO-CCR7-CD27-), confirming that T cells that are terminally differentiated and predisposed to cell senescence are infrequent. Moreover, the array of polyfunctional and cytotoxic signatures identified in the diseased airways indicated that T cells remained functionally competent.

The drivers of T-cell dynamics in the lungs are likely to be multi-factorial owing to the complex interplay between environment and host. Factors may include age-related maturation of the immune system, exposures to respiratory pathogens, iatrogenic effects, and other susceptibility determinants that dictate the inflamed milieu (14,56,57,83–86). Results of our study favor a key role for respiratory pathogens on a backdrop of immune maturation in instigating disease; these include enhanced activation of airway T cells in children who test positive for respiratory pathogens, as well as the selective accumulation of type 1-related T-cell signatures in the airways with age. Detection of RV-related T-cell signatures in the airways advances this notion. Given that all RV+ children were asymptomatic for common cold at the time of bronchoscopy, our data also imply that RV-responsive populations can remain expanded after acute infection, and are poised to exert potent effector functions. Persistence of such cells might be prolonged, as evidenced by the low frequencies of RV-related signatures identified in the airways of uninfected patients, and our ability to pinpoint “less activated” CD38- T_RM_ counterparts of RV-responsive signatures. These data echo our previous report of higher numbers of RV-specific Th1 cells in the blood of uninfected adults with asthma compared with healthy subjects, and their link to worse lung function (23). When considering host susceptibility as a driver of T-cell dynamics, our seemingly paradoxical data that correlated atopy based on serum IgE levels with higher frequencies of Th1 cells in the airways, also fits with our previous findings on virus-specific Th1 cells in adults with asthma (23). This result is notable, since it provides a plausible T-cell-based mechanism wherein the increased risk of acute wheeze in children that is known to arise from the interaction between IgE and RV fosters aberrant type 1 responses (21,25). This may also feed into T-cell plasticity and shifts in functional T-cell subsets in the atopic child, as evidenced by the inverse relationship between Th1 and Th17-like cells seen in our study.

Treatment of severe recurrent wheeze remains a major challenge. Current type 2 biologics often fail to provide symptom relief in patients with type 1 asthma. Unfortunately, biologics targeting IL-17A and its receptor have also not proven efficacious for moderate-to-severe asthma (ClinicalTrials.gov Identifiers: NCT01478360 and NCT03299686; 87,88). The complexity of T-cell types and their associated cytokines reported here, argue for the utility of combination therapy along with intervention in early life. Alternatively, identifying a single molecule that enables broad targeting of T cells might be a useful strategy. CCR5 is one such example supported by our data, whose inhibition in a mouse model of type 1 asthma has shown promise (15). That study used the CCR5 antagonist, Maraviroc, a drug currently approved to treat HIV. This drug also has the advantage of conserving CCR5+ T_RM_ that are essential to barrier immunity, while preventing egress of T cells into the lungs from the periphery (89).

Our study had several limitations. Firstly, we did not verify T-cell signatures in the tissues using endobronchial biopsies, owing to restrictions on sample procurement in children.

However, recent work in donors indicates that T cells present in the airway lumen are representative of those in the tissues (55). We also did not include age-matched healthy controls for ethical reasons. Thus, it was not possible to delineate T-cell components of barrier immunity from pathogenic T-cell types in our datasets. Nonetheless, aberrant T-cell responses in the diseased airways likely include such T-cell types or else their progeny. Regardless, our findings are substantiated by studies of airway T cells in adults that compared severe asthma with milder disease (14,17,54,59).

In summary, our results point to aberrant T-cell dynamics in the airways of children with severe recurrent wheeze. These are dominated by a complex interplay between type 1 and type 17 populations, and driven by RV-responsive elements. Our work provides the rationale for T-cell-based interventions in severe asthma of childhood in order to mitigate the vicious cycle of inflammation and corticosteroid insensitivity underlying this debilitating disease.

## Methods

### Sex as a biological variable

Our study examined both male (72%) and female (28%) patients. We did not examine sex as a biological variable.

### Human subjects

#### Pediatric

Thirty-eight children (32 for T cell phenotyping and 6 for intracellular cytokine analysis) with recurrent wheeze who underwent clinically indicated diagnostic bronchoscopies participated in the study. Children with treatment-refractory symptoms, recurrent unscheduled healthcare access for wheeze, and/or persistent airflow limitation were considered for bronchoscopy. Not all children were on ICS at the time of bronchoscopy due to either suspected structural anomalies that resulted in the suspension of corticosteroids, the inability of the family to afford the medication, or the failure of governmental health insurance providers to approve the medication. Demographic and clinical information was obtained by questionnaire and review of the medical records. Additional information on the bronchoscopy and clinical tests can be found in published reports (26,90).

#### Adult

Samples were analyzed from nine adults (age 19-26) who previously participated in a study of experimental RV-A16 infection (23,91). This included subjects with mild allergic asthma and healthy non-allergic controls. Subjects that had confounding infections or were uninfected, were excluded from analysis. Peripheral blood specimens were obtained before (day 0) and during (day 4, 7, and 21) RV infection.

#### Detection of respiratory microbes

BAL fluid was sent to the University of Virginia Medical Laboratories for bacterial cultures and multiplex-PCR for respiratory pathogens including adenovirus, coronavirus, human metapneumovirus, rhinovirus/enterovirus, Influenza A & B, Parainfluenza 1, 2, 3, & 4, respiratory syncytial virus A & B, mycoplasma pneumoniae, chlamydia pneumoniae.

#### Sample processing

PBMCs were isolated by density centrifugation using SepMate™ (Stemcell Technologies) and cryopreserved in FBS (Gibco) with 10% DMSO. Cells were isolated from BAL fluid by way of centrifugation and cryopreserved in FBS with RPMI (Gibco) and 10% DMSO.

#### T cell phenotyping spectral flow cytometry

Cells isolated from blood and BAL of 32 children with recurrent wheeze were analyzed for cell-surface and intracellular markers using a 28-color flow panel (Supplementary table 1). Blood and BAL samples were run in three batches, each containing a standard batch control sample for normalization, with matched samples being run in the same batch. In brief, cells were incubated with the viability dye LIVE/DEAD™ Fixable Blue (Invitrogen, ThermoFisher) for 15 minutes at room temperature in the dark. For detection of surface markers, cells were then washed and incubated for 40 minutes at room temperature with a 100μl mixture of antibodies, Human TruStain FcX™ (BioLegend), brilliant stain buffer plus (BD Bioscience), and FACS buffer (PBS, 0.5% bovine serum albumin, 0.1M EDTA). Cells were then washed and fixed/permeabilized (eBioscience Foxp3/Transcription Factor Staining Buffer Set, ThermoFisher) according to the manufacturer’s protocol. For intracellular markers, cells were incubated with a 100μl mixture of antibodies and permeabilization buffer for 40 minutes at room temperature. Cells were washed, resuspended in FACS buffer, and analyzed on a 5-laser Cytek® Aurora.

#### Spectral flow cytometry analysis

Following acquisition, data quality control and pre-processing was performed on all samples. This included: (1) spectral unmixing with auto-fluorescence extraction; (2) compensation; (3) manual gating to exclude doublets, dead cells, and fluorochrome aggregates; and (4) gating on the population of interest, CD3+ cells (**Supplemental Figure 12**). BAL samples with <50% viability and/or <1,000 live CD3+ cells were excluded from further analysis. Samples from the pediatric cohort were run in three batches, and to remove batch variability the normalization algorithm, CytoNorm, was applied to CD3+ cells in each sample (92). Normalization was validated through comparisons of UMAPs (93) generated pre- and post-normalization using batch control samples, as well as comparisons of histograms for each normalized marker across the batch controls (**Supplemental Figure 13**). Following data pre-processing, opt-SNE maps were generated by downsampling and pooling, (1) matched blood and BAL samples, or (2) all time points in the RV challenge for both asthmatics and controls. Intracellular markers (Ki- 67, T-bet, TCF-1) were excluded in the generation of each map. To characterize the T-cell landscape in the pediatric cohort, PhenoGraph clustering was performed using the opt-SNE axes and a K of 30. Phenograph clusters were paired down using root mean square deviation (RMSD), wherein clusters with >98% similarity were grouped. To identify populations linked to RV positivity in the BAL, the previously generated opt-SNE was analyzed by T-REX (https://github.com/cytolab/T-REX) in R (v4.4.0; Rstudio, 2024.04.2+764). To identify T-cells responding to RV infection in the challenge model, T-REX analyses were done comparing days 4, 7, or 21 of infection to day 0, separating those with asthma and healthy controls. To obtain cell frequencies for populations identified by the comparisons, T-REX files were imported into OMIQ (94), and gates were generated around populations of interest. Only populations that were not “highly variable” (no single subject comprised >50% of the cluster) were selected for further analysis. Visualization of transitions between cell populations and inference of trajectory was done using PHATE (k=20) and wishbone, on PHATE axes, in OMIQ.

#### Intracellular cytokine analysis

Cells isolated from BAL 6 children with recurrent wheeze were analyzed for surface markers and intracellular cytokines using a 21-color spectral flow cytometry panel. In brief, cells were plated in a 96-well plate and stimulated with cell activation cocktail with Brefeldin A (PMA/ionomycin, BioLegend) for 4 hours at 37°C. Following incubation, cells were harvested and incubated with the viability dye LIVE/DEAD™ Fixable Blue (Invitrogen, ThermoFisher) for 15 minutes at room temperature in the dark. For detection of surface markers, cells were then washed and incubated for 40 minutes at room temperature with a 100μl mixture of antibodies, Human TruStain FcX™ (BioLegend), brilliant stain buffer plus (BD Bioscience), and FACS buffer. Cells were then washed and fixed with BD Cytofix/Cytoperm™ Fixation/Permeabilization Kit according to manufactures protocol. For detection of intracellular cytokines, cells were permebalized with the cytofix/cytoperm kit and incubated for 30 minutes at room temperature with a 100μl mixture of antibodies, brilliant stain buffer plus, and BD Perm/Wash. Cells were then washed with BD Perm/Wash, resuspended in FACS buffer, and analyzed on a 5-laser Cytek® Aurora. Following data acquisition, quality check and pre-processing was performed as previously described. Only samples with a viability of ≥40% were included in subsequent manual gating and high-dimensional analysis. Analysis and visualization of cytokine-positive cells within T cell subsets was done using SPICE with a 1% threshold.

### Statistical analysis

Statistical analysis was performed using GraphPad Prism version 10.2.3. Differences in cell frequency of paired data were analyzed within groups by using Wilcoxon matched-pairs signed rank test or Multiple Wilcoxon test with Holm-Sidak correction for multiple comparisons. In contrast, Mann-Whitney test or Multiple Mann-Whitney with Holm-Sidak correction were used for unpaired data. Within-group comparisons over time were done using Friedman test with Dunn’s correction for multiple comparisons. Associations between T cell subset frequencies, total IgE levels, or age were determined using Spearman correlations. Adjustments for age were done using multiple logistic regressions or partial correlations. Between-group differences comparing patient characteristics were done using Kruskal-Wallis or Mann-Whitney test for continuous variables and Fisher’s exact test for categorical variables. Heatmap for PhenoGraph clusters was generated using median fluoresce intensity exported from OMIQ, and the R package ComplexHeatmap in R version 4.4.0. A p value less than 0.05 was considered significant.

### Study approval

Studies in children were approved by the University of Virginia Human Investigations Committee (Protocols #15662 and #22487). Studies in adults were approved by the University of Virginia Human Investigations Committee, and the National Institute of Allergy and Infectious Diseases Safety Committee (Protocol #12673; ClinicalTrials.gov ID: NCT02111772). Written informed consent was obtained from all adults or legal guardians, and assent was provided by children.

## Data availability

For original data, please contact. Values for data shown in graphs are reported in the Supporting Data Values file.

## Author contributions

All authors conceptualized the study. NB, LMM, and JAW devised methodology. NB performed the experiments and formal analysis. KW, WGT, and JAW provided resources. NB and JAW wrote the original draft. All authors reviewed and edited the manuscript.

## Acknowledgments

This work was supported by NIH grants (T32AI007496, R56AI178669, R21AI160334, R21AI138077, RO1AI020565, RO1AI176171-01, and UO1AI125056) and the Beirne B. Carter Center for Immunology Research.

We would like to thank Marthajoy Spano, MS, for coordinating specimen procurement. We thank Paul Dell, MS for aiding in the thawing of specimens on experiment days. We also thank Michael Solga, MS, and the staff at the University of Virginia Flow Cytometry Core Facility for their technical expertise with flow cytometry. Data generated for this publication were collected at the UVA FCCF, which is partially supported by the NCI grant (P30-CA044579). Lastly, we would like to thank all of families and patients who participated in the study.

## References

1. Center for Disease Control. Most Recent National Asthma Data. Updated May 10 2023. Accessed November 12, 2024. https://www.cdc.gov/asthma/most_recent_national_asthma_data.htm

2. Martin J, et al. Diagnosis and management of asthma in children. BMJ Paediatr Open.2022;6(1)

3. Chung KF, et al. International ERS/ATS guidelines on definition, evaluation and treatment of severe asthma. Eur Respir J.2014;43(2):343–73.

4. Chastek B, et al. Economic Burden of Illness Among Patients with Severe Asthma in a Managed Care Setting. J Manag Care Spec Pharm.2016;22(7):848–61.

5. Puranik S, et al. Predicting Severe Asthma Exacerbations in Children. Am J Respir Crit Care Med.2017;195(7):854–859.

6. Fitzpatrick AM. Severe Asthma in Children: Lessons Learned and Future Directions. J Allergy Clin Immunol Pract.2016;4(1):11–9; quiz 20-1.

7. Bacharier LB, Jackson DJ. Biologics in the treatment of asthma in children and adolescents. J Allergy Clin Immunol.2023;151(3):581–589.

8. Khurana S, et al. Management of severe asthma: summary of the European Respiratory Society/American Thoracic Society task force report. Breathe (Sheff).2020;16(2):200058.

9. Fahy JV. Type 2 inflammation in asthma--present in most, absent in many. Nat Rev Immunol.2015;15(1):57–65.

10. Harker JA, Lloyd CM. T helper 2 cells in asthma. J Exp Med.2023;220(6)

11. Papadopoulos NG, et al. Type 2 Inflammation and Asthma in Children: A Narrative Review. J Allergy Clin Immunol Pract.2024;12(9):2310–2324.

12. Romeo MJ, et al. A molecular perspective on TH2-promoting cytokine receptors in patients with allergic disease. J Allergy Clin Immunol.2014;133(4):952–60.

13. Woodfolk JA. T-cell responses to allergens. J Allergy Clin Immunol.2007;119(2):280–94; quiz 295-6.

14. Camiolo MJ, et al. High-dimensional profiling clusters asthma severity by lymphoid and non- lymphoid status. Cell Rep.2021;35(2):108974.

15. Gauthier M, et al. Dual role for CXCR3 and CCR5 in asthmatic type 1 inflammation. J Allergy Clin Immunol.2022;149(1):113–124.e7.

16. Gauthier M, et al. CCL5 is a potential bridge between type 1 and type 2 inflammation in asthma. J Allergy Clin Immunol.2023;152(1):94–106.e12.

17. Herrera-De La Mata S, et al. Cytotoxic CD4(+) tissue-resident memory T cells are associated with asthma severity. Med. 2023;4(12):875-897.e8.

18. Oriss TB, et al. IRF5 distinguishes severe asthma in humans and drives Th1 phenotype and airway hyperreactivity in mice. JCI Insight.2017;2(10)

19. Wisniewski JA, et al. T(H)1 signatures are present in the lower airways of children with severe asthma, regardless of allergic status. J Allergy Clin Immunol.2018;141(6):2048-2060.e13.

20. Hansel TT, et al. A Comprehensive Evaluation of Nasal and Bronchial Cytokines and Chemokines Following Experimental Rhinovirus Infection in Allergic Asthma: Increased Interferons (IFN-γ and IFN-λ) and Type 2 Inflammation (IL-5 and IL-13). EBioMedicine.2017;19:128–138.

21. Heymann PW, et al. Viral infections in relation to age, atopy, and season of admission among children hospitalized for wheezing. J Allergy Clin Immunol.2004;114(2):239–47.

22. Johnston SL, et al. Community study of role of viral infections in exacerbations of asthma in 9-11 year old children. Bmj.1995;310(6989):1225-9.

23. Muehling LM, et al. Human T(H)1 and T(H)2 cells targeting rhinovirus and allergen coordinately promote allergic asthma. J Allergy Clin Immunol.2020;146(3):555–570.

24. Schwantes EA, et al. Interferon gene expression in sputum cells correlates with the Asthma Index Score during virus-induced exacerbations. Clin Exp Allergy.2014;44(6):813–21.

25. Soto-Quiros M, et al. High titers of IgE antibody to dust mite allergen and risk for wheezing among asthmatic children infected with rhinovirus. J Allergy Clin Immunol.2012;129(6):1499–1505.e5.

26. Teague WG, et al. A novel syndrome of silent rhinovirus-associated bronchoalveolitis in children with recurrent wheeze. J Allergy Clin Immunol.2024;

27. Khetsuriani N, et al. Prevalence of viral respiratory tract infections in children with asthma. J Allergy Clin Immunol.2007;119(2):314–21.

28. Jartti T, et al. Serial viral infections in infants with recurrent respiratory illnesses. Eur Respir J.2008;32(2):314–20.

29. Gauthier M, et al. Severe asthma in humans and mouse model suggests a CXCL10 signature underlies corticosteroid-resistant Th1 bias. JCI Insight.2017;2(13)

30. Arnoux B, et al. Increased bronchoalveolar lavage CD8 lymphocyte subset population in wheezy infants. Pediatr Allergy Immunol.2001;12(4):194–200.

31. Marguet C, et al. Bronchoalveolar cell profiles in children with asthma, infantile wheeze, chronic cough, or cystic fibrosis. Am J Respir Crit Care Med.1999;159(5 Pt 1):1533-40.

32. Kohlmeier JE, et al. The chemokine receptor CCR5 plays a key role in the early memory CD8+ T cell response to respiratory virus infections. Immunity.2008;29(1):101–13.

33. Galkina E, et al. Preferential migration of effector CD8+ T cells into the interstitium of the normal lung. J Clin Invest.2005;115(12):3473–83.

34. Brinkmann V, Kristofic C. Regulation by corticosteroids of Th1 and Th2 cytokine production in human CD4+ effector T cells generated from CD45RO- and CD45RO+ subsets. J Immunol.1995;155(7):3322–8.

35. Gemou-Engesaeth V, et al. Inhaled glucocorticoid therapy of childhood asthma is associated with reduced peripheral blood T cell activation and ’Th2-type’ cytokine mRNA expression. Pediatrics.1997;99(5):695–703.

36. Kaur M, et al. The effects of corticosteroids on cytokine production from asthma lung lymphocytes. Int Immunopharmacol.2014;23(2):581–4.

37. Levine JH, et al. Data-Driven Phenotypic Dissection of AML Reveals Progenitor-like Cells that Correlate with Prognosis. Cell.2015;162(1):184–97.

38. Mackay LK, et al. T-box Transcription Factors Combine with the Cytokines TGF-β and IL-15 to Control Tissue-Resident Memory T Cell Fate. Immunity.2015;43(6):1101–11.

39. Oja AE, et al. Trigger-happy resident memory CD4(+) T cells inhabit the human lungs. Mucosal Immunol.2018;11(3):654–667.

40. Basdeo SA, et al. Polyfunctional, Pathogenic CD161+ Th17 Lineage Cells Are Resistant to Regulatory T Cell-Mediated Suppression in the Context of Autoimmunity. J Immunol.2015;195(2):528–40.

41. Basdeo SA, et al. Ex-Th17 (Nonclassical Th1) Cells Are Functionally Distinct from Classical Th1 and Th17 Cells and Are Not Constrained by Regulatory T Cells. J Immunol.2017;198(6):2249–2259.

42. Escobar G, et al. T cell factor 1: A master regulator of the T cell response in disease. Sci Immunol.2020;5(53)

43. Wu J, et al. T Cell Factor 1 Suppresses CD103+ Lung Tissue-Resident Memory T Cell Development. Cell Rep.2020;31(1):107484.

44. Moon KR, et al. Visualizing structure and transitions in high-dimensional biological data. Nat Biotechnol.2019;37(12):1482–1492.

45. Kumar BV, et al. Human Tissue-Resident Memory T Cells Are Defined by Core Transcriptional and Functional Signatures in Lymphoid and Mucosal Sites. Cell Rep.2017;20(12):2921–2934.

46. DeRogatis JM, et al. Cell-Intrinsic CD38 Expression Sustains Exhausted CD8(+) T Cells by Regulating Their Survival and Metabolism during Chronic Viral Infection. J Virol.2023;97(4):e0022523.

47. De Simone G, et al. CXCR3 Identifies Human Naive CD8(+) T Cells with Enhanced Effector Differentiation Potential. J Immunol.2019;203(12):3179–3189.

48. Barone SM, et al. Unsupervised machine learning reveals key immune cell subsets in COVID-19, rhinovirus infection, and cancer therapy. Elife.2021;10

49. Teague WG, et al. Novel Treatment-Refractory Preschool Wheeze Phenotypes Identified by Cluster Analysis of Lung Lavage Constituents. J Allergy Clin Immunol Pract.2021;9(7):2792–2801.e4.

50. Kloepfer KM, et al. Detection of pathogenic bacteria during rhinovirus infection is associated with increased respiratory symptoms and asthma exacerbations. J Allergy Clin Immunol.2014;133(5):1301–7, 1307.e1-3.

51. Diggins KE, et al. Characterizing cell subsets using marker enrichment modeling. Nat Methods.2017;14(3):275–278.

52. Roederer M, et al. SPICE: exploration and analysis of post-cytometric complex multivariate datasets. Cytometry A.2011;79(2):167–74.

53. Wacleche VS, et al. Identification of T Peripheral Helper (Tph) Cells. Methods Mol Biol.2022;2380:59–76.

54. Raundhal M, et al. High IFN-γ and low SLPI mark severe asthma in mice and humans. J Clin Invest.2015;125(8):3037–50.

55. Snyder ME, et al. Generation and persistence of human tissue-resident memory T cells in lung transplantation. Sci Immunol.2019;4(33)

56. Connors TJ, et al. Site-specific development and progressive maturation of human tissue-resident memory T cells over infancy and childhood. Immunity.2023;56(8):1894–1909.e5.

57. Thome JJ, et al. Spatial map of human T cell compartmentalization and maintenance over decades of life. Cell.2014;159(4):814–28.

58. Al-Ramli W, et al. T(H)17-associated cytokines (IL-17A and IL-17F) in severe asthma. J Allergy Clin Immunol.2009;123(5):1185–7.

59. Doe C, et al. Expression of the T helper 17-associated cytokines IL-17A and IL-17F in asthma and COPD. Chest.2010;138(5):1140–7.

60. Chien JW, et al. Increased IL-17A secreting CD4+ T cells, serum IL-17 levels and exhaled nitric oxide are correlated with childhood asthma severity. Clin Exp Allergy.2013;43(9):1018–26.

61. Molet S, et al. IL-17 is increased in asthmatic airways and induces human bronchial fibroblasts to produce cytokines. J Allergy Clin Immunol.2001;108(3):430–8.

62. Chang Y, et al. TH17 cytokines induce human airway smooth muscle cell migration. J Allergy Clin Immunol.2011;127(4):1046–53.e1-2.

63. Chang Y, et al. Th17-associated cytokines promote human airway smooth muscle cell proliferation. Faseb j.2012;26(12):5152–60.

64. Celada LJ, et al. PD-1 up-regulation on CD4(+) T cells promotes pulmonary fibrosis through STAT3- mediated IL-17A and TGF-β1 production. Sci Transl Med.2018;10(460)

65. Chen Y, et al. Stimulation of airway mucin gene expression by interleukin (IL)-17 through IL-6 paracrine/autocrine loop. J Biol Chem.2003;278(19):17036–43.

66. Kudo M, et al. IL-17A produced by αβ T cells drives airway hyper-responsiveness in mice and enhances mouse and human airway smooth muscle contraction. Nat Med.2012;18(4):547–54.

67. Laan M, et al. Neutrophil recruitment by human IL-17 via C-X-C chemokine release in the airways. J Immunol.1999;162(4):2347–52.

68. Banuelos J, et al. BCL-2 protects human and mouse Th17 cells from glucocorticoid-induced apoptosis. Allergy.2016;71(5):640–50.

69. de Castro Kroner J, et al. Glucocorticoids promote intrinsic human T(H)17 differentiation. J Allergy Clin Immunol.2018;142(5):1669–1673.e11.

70. Vazquez-Tello A, et al. Induction of glucocorticoid receptor-beta expression in epithelial cells of asthmatic airways by T-helper type 17 cytokines. Clin Exp Allergy.2010;40(9):1312–22.

71. Gray JI, et al. Human γδ T cells in diverse tissues exhibit site-specific maturation dynamics across the life span. Sci Immunol.2024;9(96):eadn3954.

72. Kang I, et al. Double-edged sword: γδ T cells in mucosal homeostasis and disease. Exp Mol Med.2023;55(9):1895–1904.

73. Salter B, et al. Airway autoantibodies are determinants of asthma severity. Eur Respir J.2022;60(6)

74. Son K, et al. Autoantibody-mediated Macrophage Dysfunction in Patients with Severe Asthma with Airway Infections. Am J Respir Crit Care Med.2023;207(4):427–437.

75. Shivakumar S, et al. T cell receptor alpha/beta expressing double-negative (CD4-/CD8-) and CD4+ T helper cells in humans augment the production of pathogenic anti-DNA autoantibodies associated with lupus nephritis. J Immunol.1989;143(1):103–12.

76. Crispín JC, et al. Expanded double negative T cells in patients with systemic lupus erythematosus produce IL-17 and infiltrate the kidneys. J Immunol.2008;181(12):8761–6.

77. Kleinschek MA, et al. Circulating and gut-resident human Th17 cells express CD161 and promote intestinal inflammation. J Exp Med.2009;206(3):525–34.

78. Yokoi T, et al. Identification of a unique subset of tissue-resident memory CD4(+) T cells in Crohn’s disease. Proc Natl Acad Sci U S A.2023;120(1):e2204269120.

79. Peng C, et al. Engagement of the costimulatory molecule ICOS in tissues promotes establishment of CD8(+) tissue-resident memory T cells. Immunity.2022;55(1):98–114.e5.

80. Tong J, et al. Fas-positive T cells regulate the resolution of airway inflammation in a murine model of asthma. J Exp Med.2006;203(5):1173–84.

81. Weisberg SP, et al. Tissue-Resident Memory T Cells Mediate Immune Homeostasis in the Human Pancreas through the PD-1/PD-L1 Pathway. Cell Rep.2019;29(12):3916–3932.e5.

82. Bouillet P, O’Reilly LA. CD95, BIM and T cell homeostasis. Nat Rev Immunol.2009;9(7):514–9.

83. Connors TJ, et al. Airway CD8(+) T Cells Are Associated with Lung Injury during Infant Viral Respiratory Tract Infection. Am J Respir Cell Mol Biol.2016;54(6):822-30.

84. Connors TJ, et al. Developmental Regulation of Effector and Resident Memory T Cell Generation during Pediatric Viral Respiratory Tract Infection. J Immunol.2018;201(2):432–439.

85. Poon MML, et al. Heterogeneity of human anti-viral immunity shaped by virus, tissue, age, and sex. Cell Rep.2021;37(9):110071.

86. Shanthikumar S, et al. Mapping Pulmonary and Systemic Inflammation in Preschool Aged Children With Cystic Fibrosis. Front Immunol.2021;12:733217.

87. Bryant N, Muehling LM. T-cell responses in asthma exacerbations. Ann Allergy Asthma Immunol.2022;129(6):709–718.

88. Busse WW, et al. Randomized, double-blind, placebo-controlled study of brodalumab, a human anti- IL-17 receptor monoclonal antibody, in moderate to severe asthma. Am J Respir Crit Care Med.2013;188(11):1294–302.

89. Woodward Davis AS, et al. The human tissue-resident CCR5(+) T cell compartment maintains protective and functional properties during inflammation. Sci Transl Med.2019;11(521)

90. Teague WG, et al. Lung Lavage Granulocyte Patterns and Clinical Phenotypes in Children with Severe, Therapy-Resistant Asthma. J Allergy Clin Immunol Pract.2019;7(6):1803–1812.e10.

91. Heymann PW, et al. Understanding the asthmatic response to an experimental rhinovirus infection: Exploring the effects of blocking IgE. J Allergy Clin Immunol.2020;146(3):545–554.

92. Van Gassen S, et al. CytoNorm: A Normalization Algorithm for Cytometry Data. Cytometry A.2020;97(3):268–278.

93. Becht E, et al. Dimensionality reduction for visualizing single-cell data using UMAP. Nature Biotechnology.2019;37(1):38-+.

94. Dotmatics. OMIQ. Accessed November 12, 2024. www.omiq.ai

