## Supplemental Figures for "Rhinovirus as a Driver of Airway T-Cell Dynamics in Children with Severe Asthma"

Supplemental table 1. Spectral flow cytometry panel for T cell phenotyping.

| Marker | Color | Clone | Vendor | Catalog number | Dilution |
|---|---|---|---|---|---|
| Viability | Live/Dead Blue | | Invitrogen | L23105 | 1:4000 |
| CD3 | BUV395 | SK7 | BD | 564001 | 1:50 |
| CD4 | BUV496 | OKT4 | BD | 750980 | 1:50 |
| CD45RO | BUV563 | UCHL1 | BD | 748369 | 1:100 |
| PD-1 | BUV615 | EH12.1 | BD | 612991 | 1:50 |
| CXCR3 | BUV661 | 1C6/CXCR3 | BD | 741649 | 1:50 |
| CCR5 | BUV737 | 2D7/CCR5 | BD | 612808 | 1:25 |
| IL-4Rα | BV421 | G077F6 | Biolegend | 355014 | 1:50 |
| CRTH2 | BV480 | BM16 | BD | 746388 | 1:200 |
| CCR4 | BV605 | L291H4 | Biolegend | 359418 | 1:25 |
| CCR7 | BV650 | G043H7 | Biolegend | 353234 | 1:20 |
| CCR6 | BV711 | G034E3 | Biolegend | 353436 | 1:50 |
| CXCR5 | BV750 | J252D4 | Biolegend | 356942 | 1:50 |
| CD161 | BV785 | HP-3G10 | Biolegend | 339930 | 1:20 |
| ICOS | BB515 | C398.4A | BD | 565881 | 1:100 |
| CD14 | FITC | M5E2 | Biolegend | 301804 | 1:20 |
| CD19 | FITC | HIB19 | Biolegend | 302206 | 1:20 |
| CD8 | PerCP | SK1 | Biolegend | 344708 | 1:200 |
| CD95 | BB700 | DX2 | BD | 566543 | 1:100 |
| CD103 | Per-CP eFluor710 | Ber-ACT8 | Invitrogen | 46103742 | 1:50 |
| ST2 | PE | Polyclonal | R&D Systems | FAB5231P100 | 1:10 |
| CD25 | PE-FIRE700 | M-A251 | Biolegend | 356146 | 1:50 |
| T-bet | PE-CY7 | 4B10 | Biolegend | 644823 | 1:800 |
| Ki-67 | APC | 20Raj1 | Invitrogen | 17-5699-41 | 1:200 |
| TCF-1 | AF647 | C63D9 | Cell Signaling Technology | 6709S | 1:200 |
| CD127 | APC-R700 | HIL-7R-M21 | BD | 565185 | 1:200 |
| CD27 | APC-CY7 | O323 | Biolegend | 302816 | 1:50 |
| CD38 | APC-FIRE810 | HIT2 | Biolegend | 303550 | 1:50 |

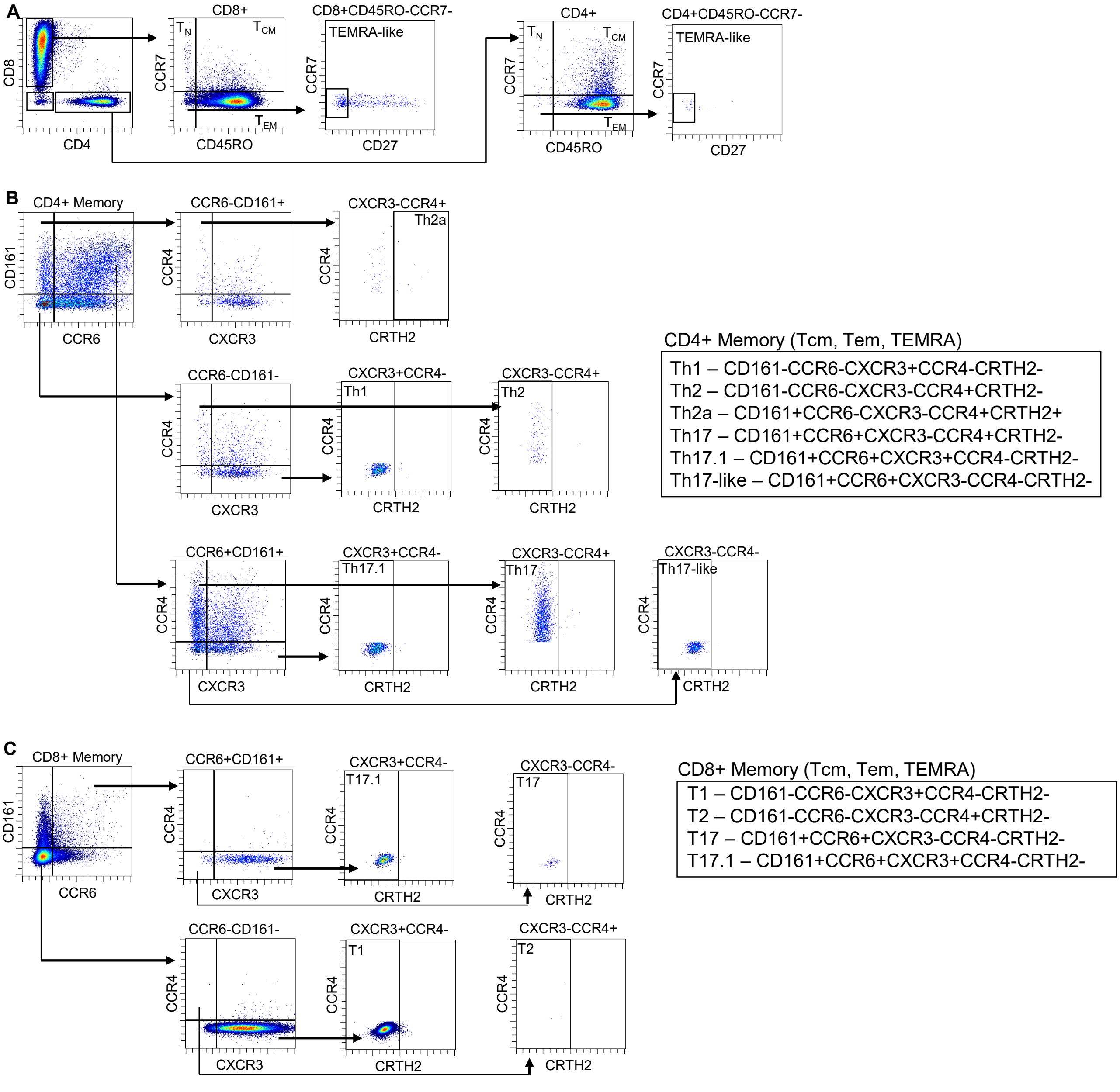


**Supplemental Figure 1.** Representative scatter plots showing gating strategy for (A) memory T cells and (B and C) T cell subsets.

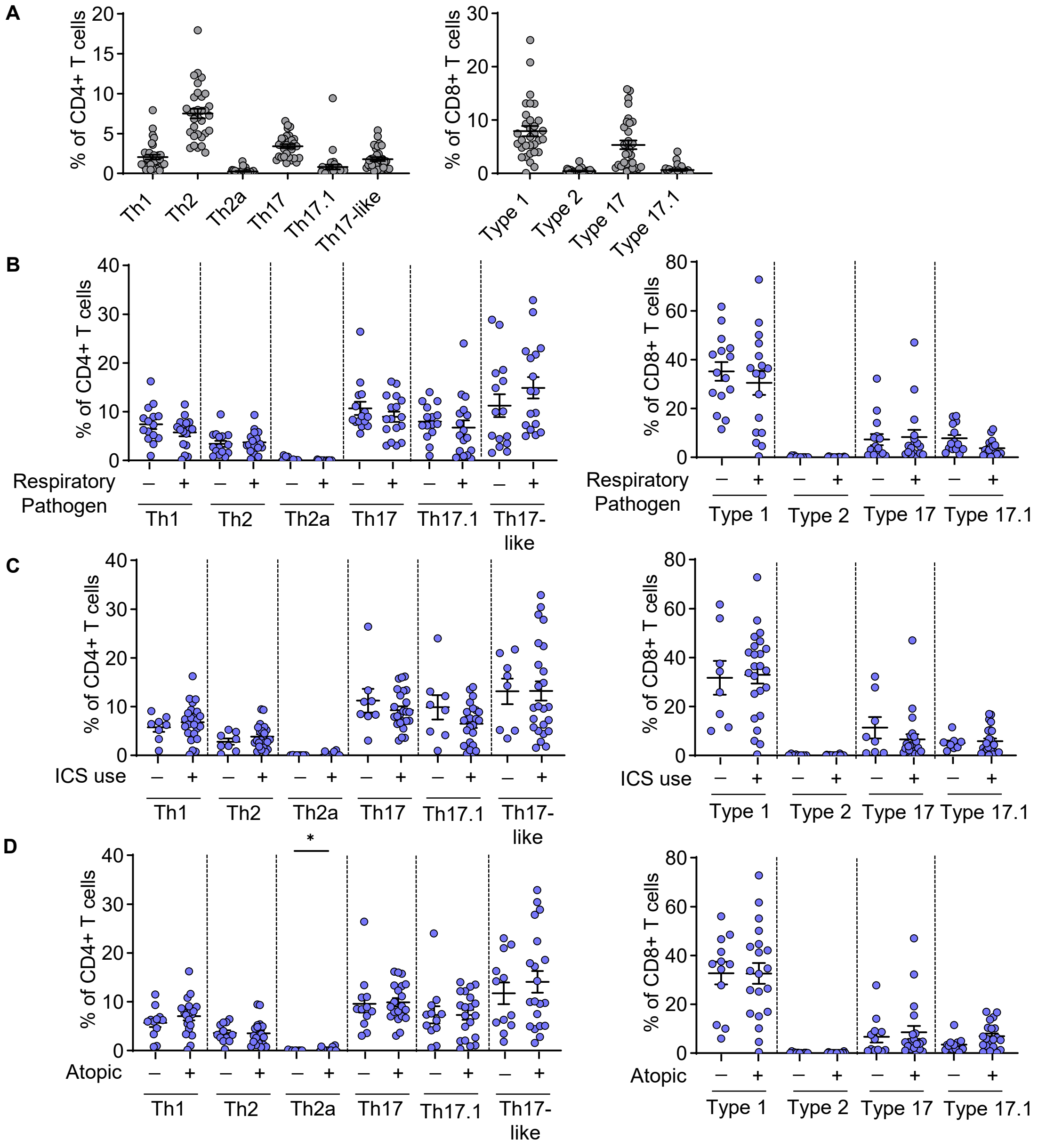


**Supplemental Figure 2. The influence of clinical characteristics on T cell subsets in the airways.** (A) T cell subset frequency in the blood (n=32). T cell subset frequency in the BAL, delineated by (B) the presence of respiratory pathogens (- n=15, + n=17), (C) inhaled corticosteroid (ICS) use (- n=8, + n=24), and (D) atopy (- n=12, + n=20) (D) in BAL. Multiple Mann-Whitney tests with Holm-Sidak correction was used to calculate significance (B-D). *p<0.05

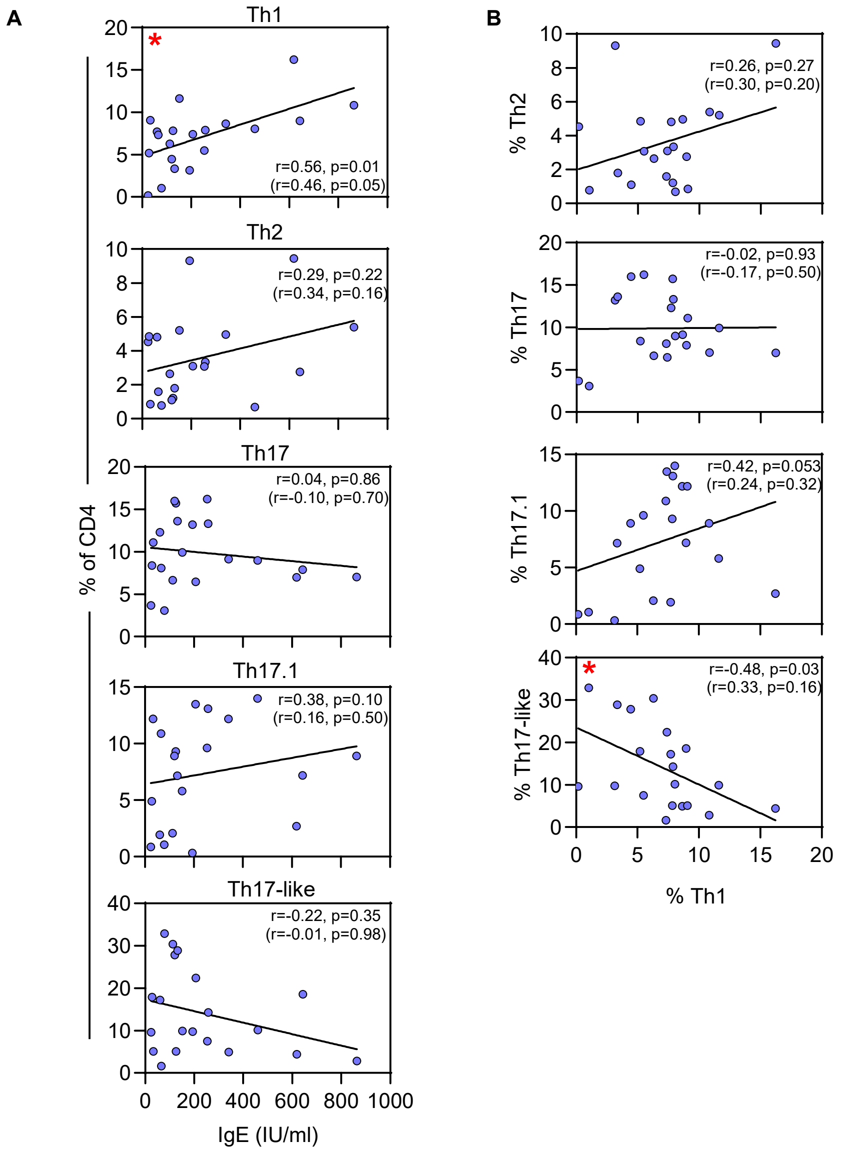


**Supplemental Figure 3. Th1 cells are linked to total IgE in atopic patients.** Spearman correlations for (A) CD4+ T cell subset frequency vs. IgE levels and (B) Th1 cell frequency vs. other CD4+ subsets, for atopic patients only (n=20). Correlation values in the parentheses were corrected for age. Line denotes linear regression. Red asterisk denotes significant correlations.

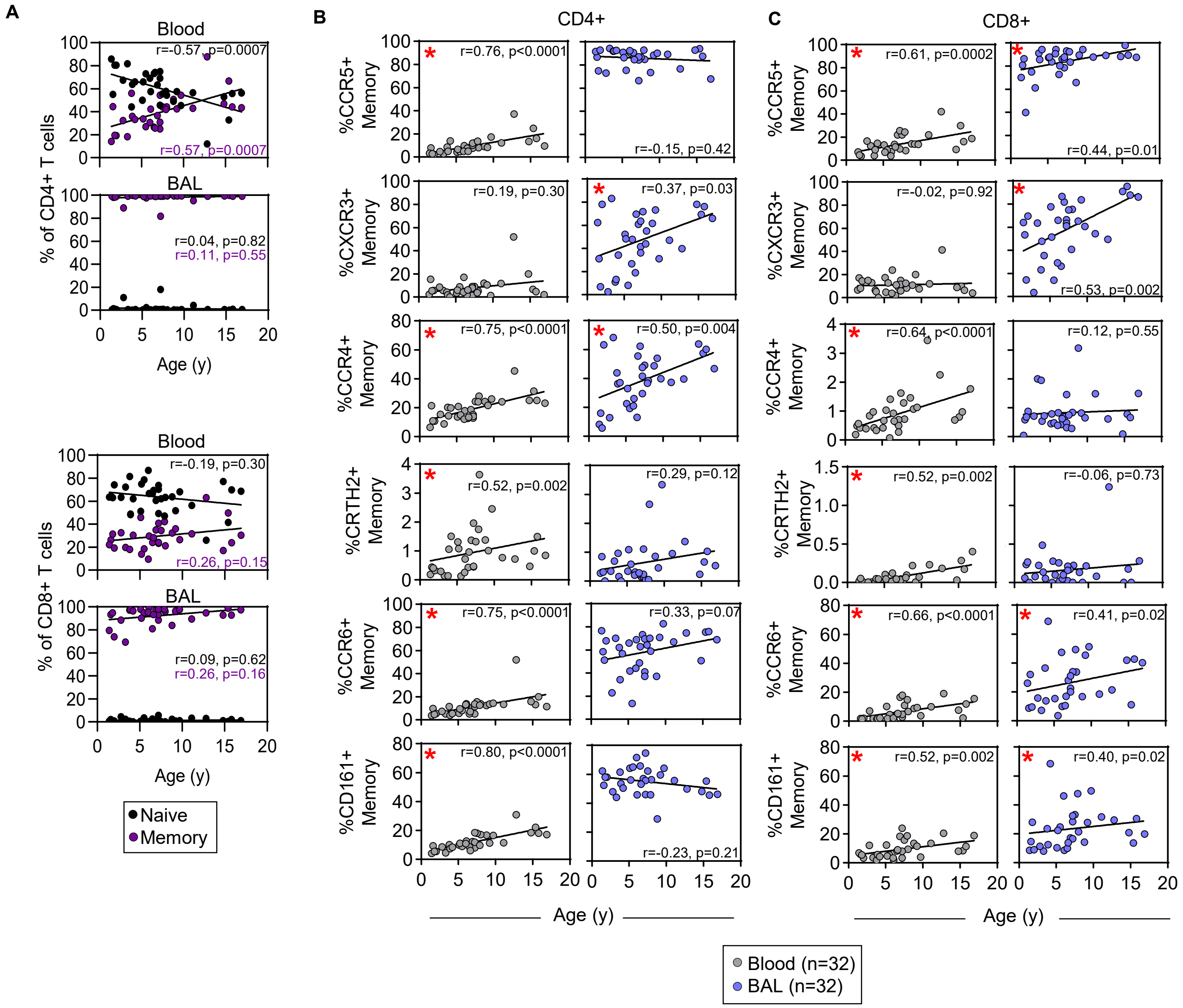


**Supplemental Figure 4. Preferential accumulation of type 1 T cells in the airways with age.** (A) Spearman correlations for naïve and memory ($T_{CM}$, $T_{EM}$, TEMRA-like) CD4+ and CD8+ T cells vs. age in blood and BAL (n=32). Spearman correlations for receptor-positive memory (B) CD4+ and (C) CD8+ T cells vs. age (years). Line denotes linear regression. Red asterisks denote significant correlations (B and C).

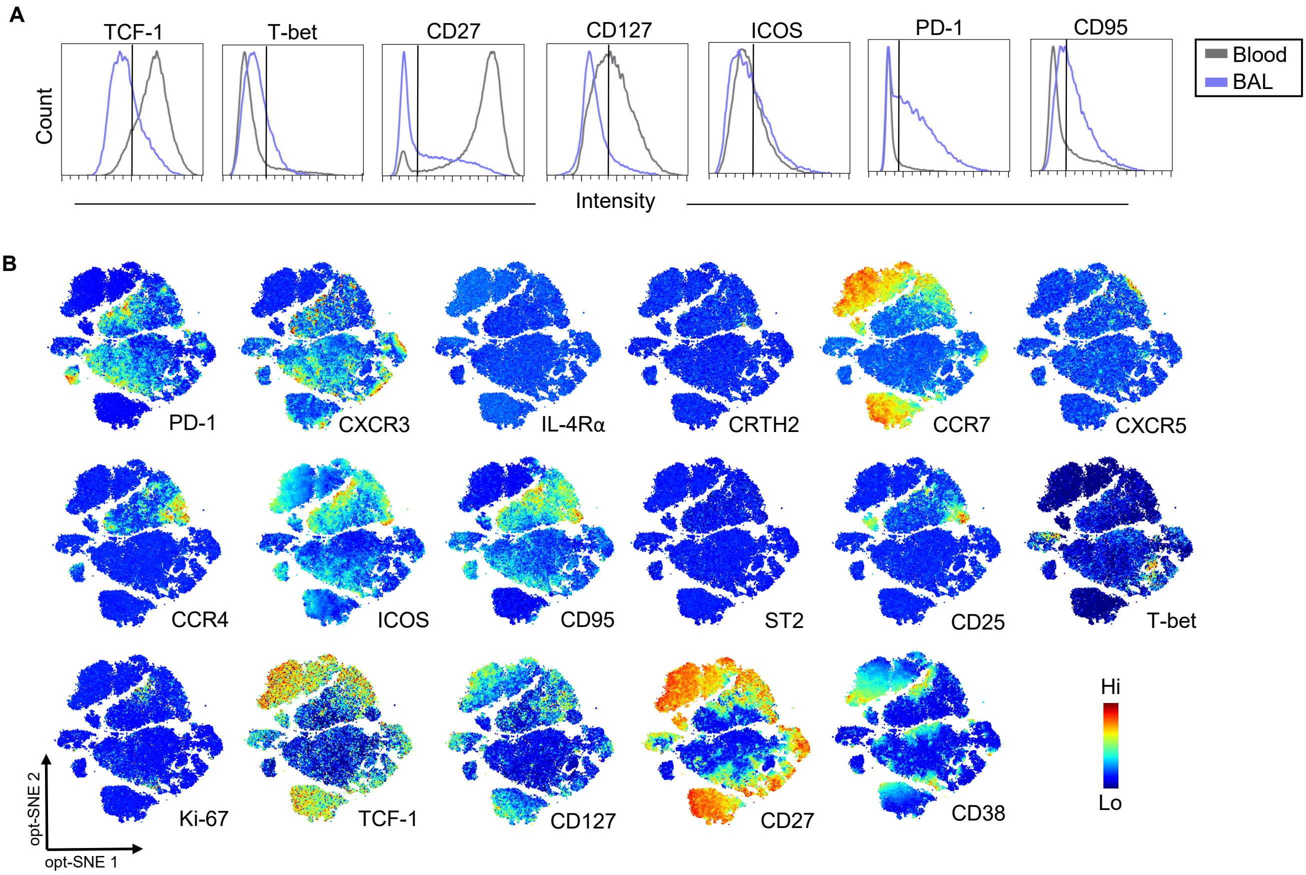


**Supplemental Figure 5. T-cell landscape in children with recurrent wheeze.** (A) Histograms showing median expression of select markers on blood or BAL CD3+ cells. (B) Heatmaps showing the distribution of marker expression on opt-SNE axes.

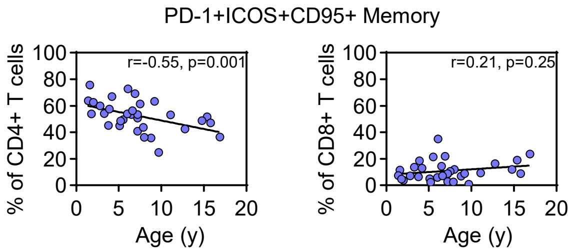


**Supplemental Figure 6. T-cell activation in the lower airways changes with age.** Spearman correlation of activated (PD-1+ICOS+CD95+) CD4+ or CD8+ memory T cells vs. age (years) (n=32). Line denotes linear regression.

Supplemental Table 2. Demographic and clinical data comparing RV- and RV+ subjects

| | **RV- (n=22)** | **RV+ (n=10)** | **P value[D]** |
|---|---|---|---|
| Age (years)[A] | 6.7 (5.0-8.9) | 4.8 (3.0-7.7) | 0.19[E] |
| Male sex, n (%) | 16 (72.7) | 7 (70.0) | >0.99[F] |
| White, n (%) | 16 (72.7) | 8 (80.0) | >0.99[F] |
| Total IgE (IU/ml)[A] | 72.7 (33.9-156.0) | 43.5 (14.8-128.1) | 0.35[F] |
| Atopic, n (%) | 15 (68.2) | 5 (50.0) | 0.44[I] |
| Medication, n (%) | | | |
| Inhaled corticosteroid (ICS) | 16 (72.7) | 8 (80.0) | >0.99[F] |
| Long-acting β-agonist | 13 (59.1) | 1 (10.0) | 0.02[F] |
| Antileukotriene | 7 (31.9) | 2 (20.0) | 0.68[F] |
| Oral prednisone | 2 (9.1) | 1 (10.0) | >0.99[F] |
| Dupilumab | 2 (9.1) | 0 (0.0) | >0.99[F] |
| Mepolizumab | 1 (4.5) | 0 (0.0) | >0.99[F] |
| Omalizumab | 1 (4.5) | 0 (0.0) | >0.99[F] |
| BAL composition %[B] | (n=22) | (n=9) | |
| Eosinophils | 0% (0-4) | 0% (0-14) | 0.43[E] |
| Neutrophils | 2.5% (0-68) | 29% (0-90) | 0.03[E] |
| Macrophage | 75.5% (0-95) | 47% (0-88) | 0.05[E] |
| Lymphocytes | 4% (0-11) | 3% (0-9) | 0.60[E] |
| Infection status, n (%) | | | |
| Rhinovirus | 0 (0.0) | 10 (100.0) | <0.0001[F] |
| Other virus[C] | 3 (13.6) | 0 (0.0) | 0.53[F] |
| Any bacteria | 6 (27.3) | 4 (40.0) | 0.68[F] |
| Co-infection | 2 (9.1) | 4 (40.0) | 0.06[F] |

[A] Geometric mean (95% CI), [B] Median (range), [C] Adenovirus and Metapneumovirus, [D] RV- v RV+, [E] Mann-Whitney test, [F] Fisher's exact test.

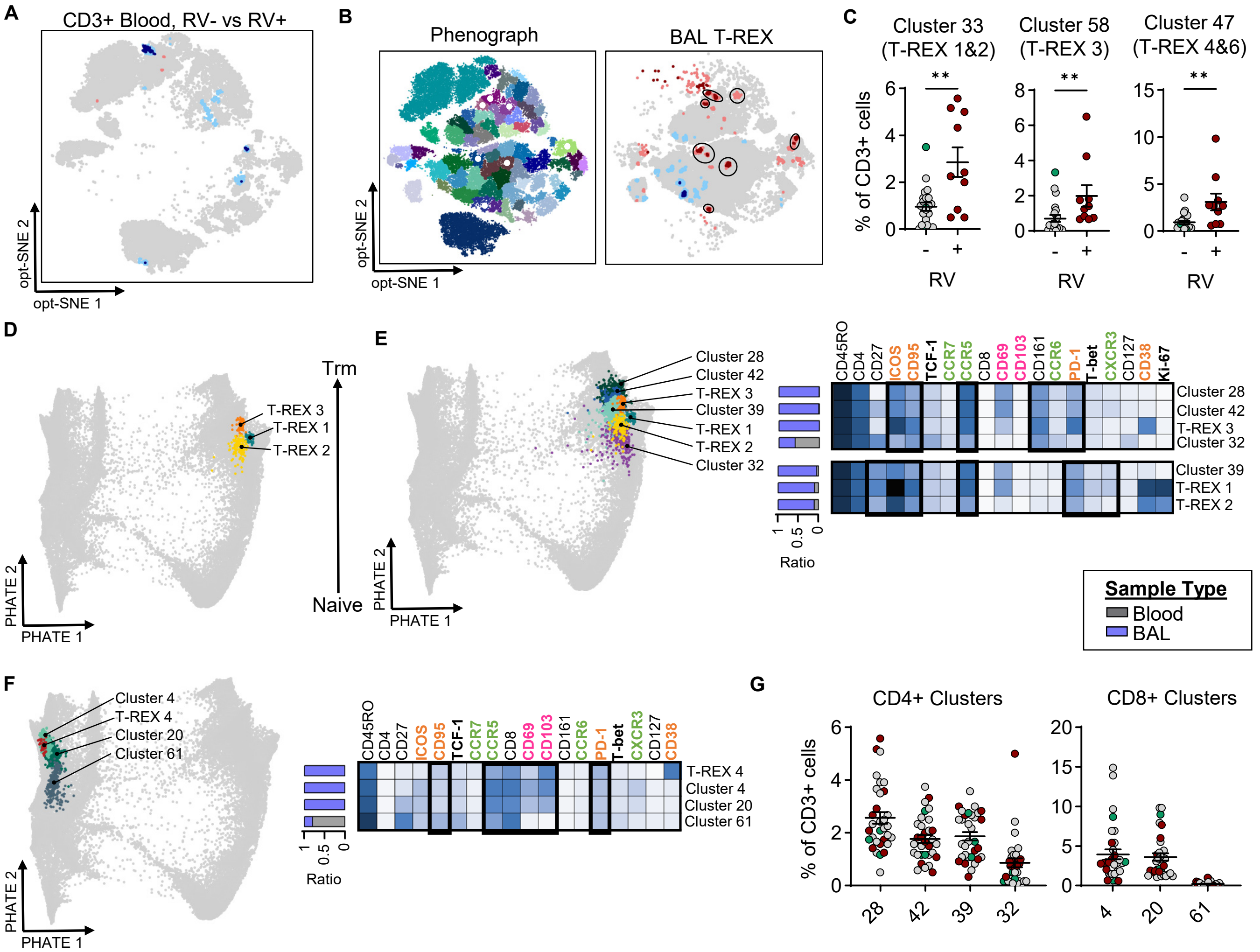

**Supplemental Figure 7. RV-related signatures persist in the airways.** (A) T-REX comparison of T cells in the blood, between RV- (n=22) and RV+ (n=10) patients. (B) T-REX populations in relation to PhenoGraph clusters. White dots on left opt-SNE show the PhenoGraph clusters that correspond to T-REX populations. (C) Frequency of corresponding PhenoGraph clusters in RV- and RV+ patients. (D) CD4+ T-REX populations (1-3) projected on PHATE map. (E and F) PHATE map showing PhenoGraph clusters related to T-REX populations. Heatmaps comparing marker expression between PhenoGraph clusters and T-REX populations. Black squares on the heatmap denote markers that are similar between the populations. Annotations on the left denote the proportion of cells with each cluster that come from blood or BAL. (G) Frequency of PhenoGraph clusters from panels E and F, in the BAL. For data points in C and G, green are patients positive for other viruses (n=3), red are those positive for RV (n=10), and grey are those negative for any virus (=19). Horizontal bars denote mean ± SEM. Mann-Whitney test was used to calculate significance (C). **p≤0.01

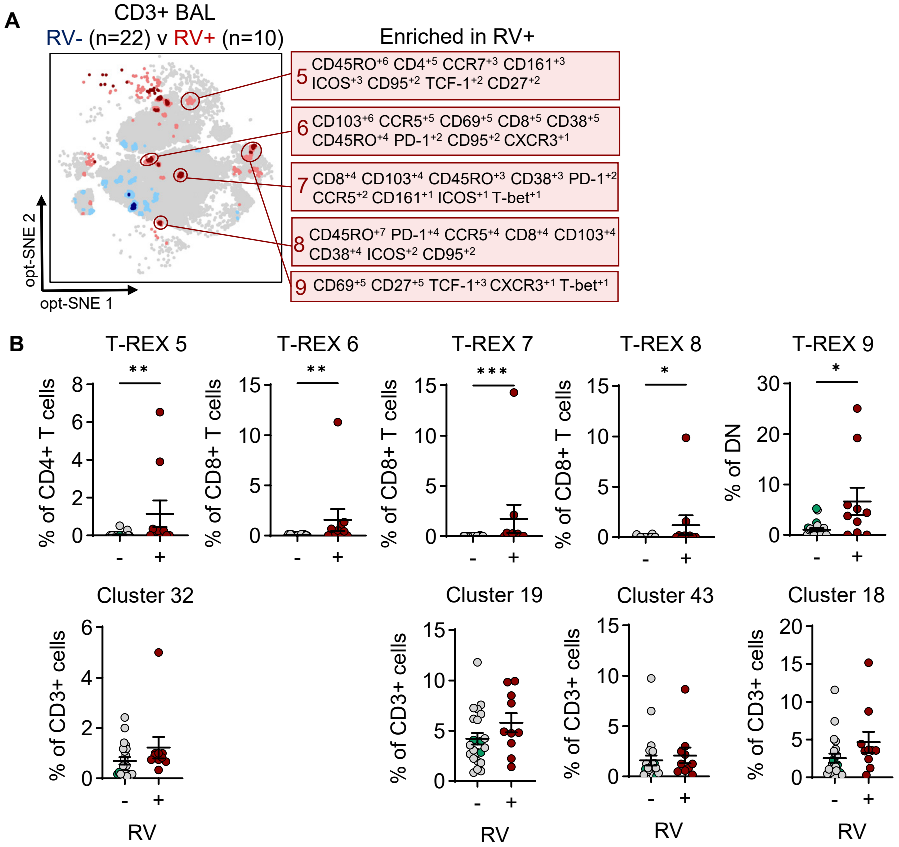


**Supplemental Figure 8. “Highly variable” RV-related T cell signatures.** (A) MEM signatures of “highly variable” populations enriched in the RV+ group. (B) Frequency of “highly variable” T-REX populations enriched in RV+ group and corresponding PhenoGraph clusters. The corresponding cluster for T-REX 6 was shown in supplemental figure 8C. Green data points in RV- group denote subjects positive for other viruses (n=3). Horizontal bars denote mean ± SEM. Mann-Whitney test used to calculate significance (B). *p<0.05, **p≤0.01, ***p≤0.001.

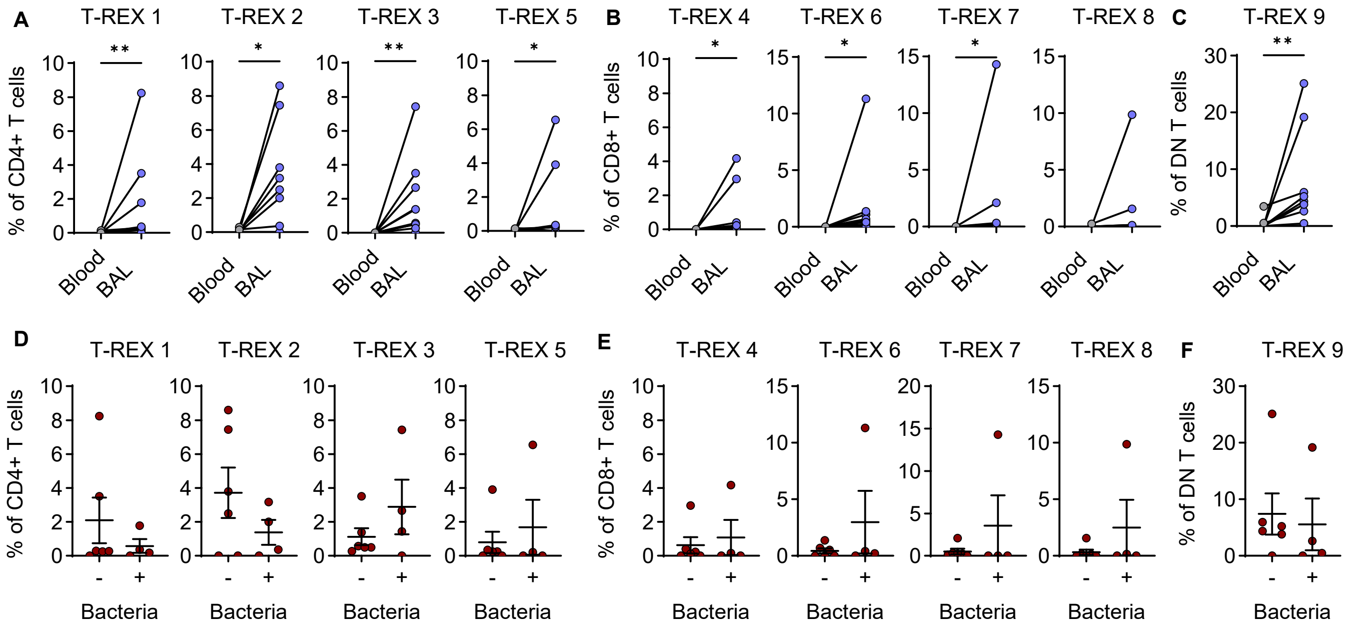


**Supplemental Figure 9. RV-related signatures are enriched in the lower airways compared to blood.** Comparison of (A) CD4+, (B) CD8+, and (C) DN T-REX populations in blood and BAL. Frequency of (D) CD4+, (E) CD8+, and (F) DN T-REX populations delineated by the presence (n=4) or absence (n=6) of bacteria. Data shown is only for RV+ patients (n=10). Lines denote mean ± SEM. Wilcoxon matched-pairs signed rank test (A-C) and Mann-Whitney test (D-F) used to calculate significance. *p<0.05, **p≤0.01.

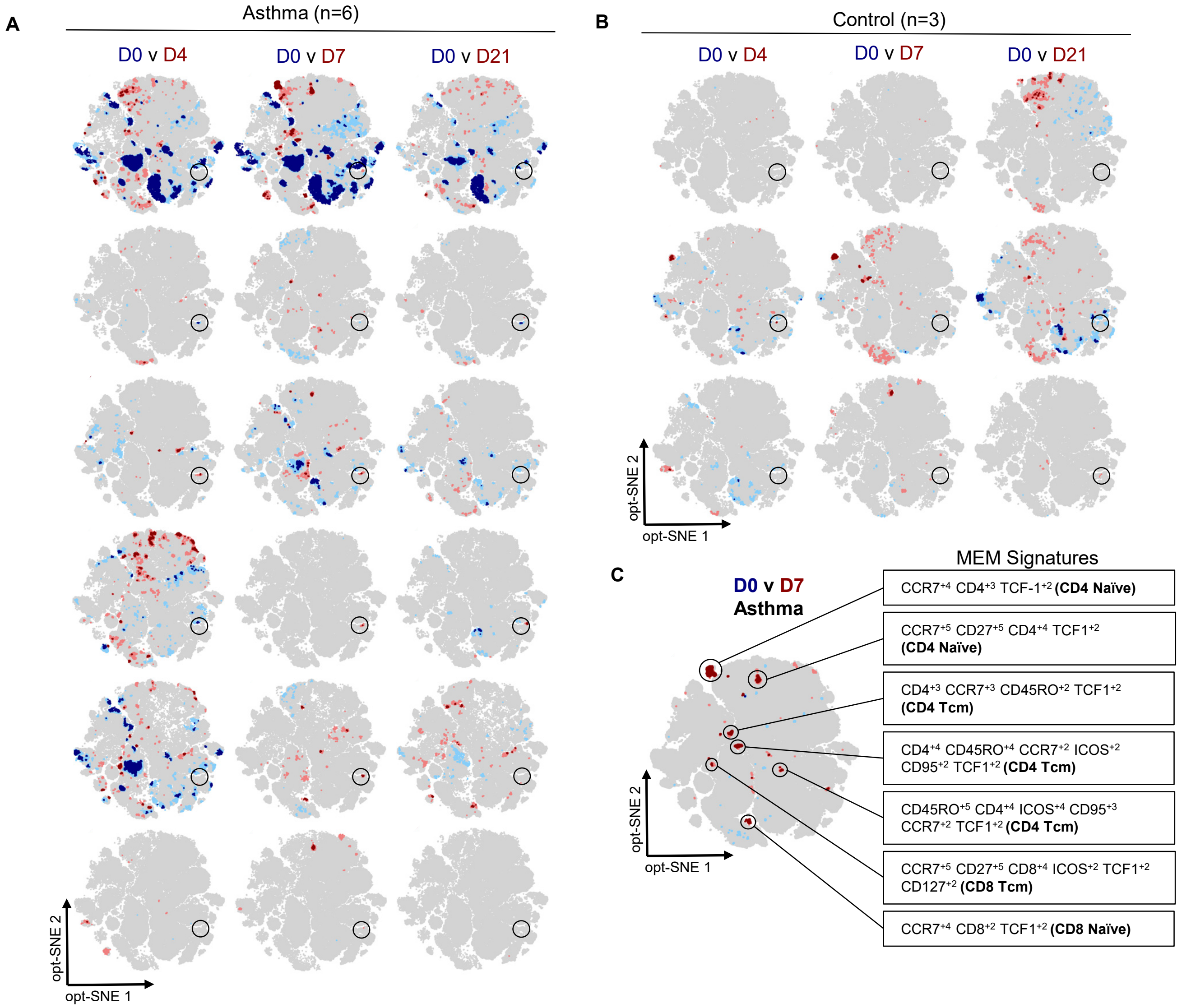


**Supplemental Figure 10. T-cell responses to experimental RV infection are heterogeneous.** Individual T-REX comparisons of days 4, 7, and 21 of infection to day 0, for (A) adults with allergic asthma and (B) healthy controls. Black circles denote the location of T-REX A from Figure 6C. (C) MEM signatures of “highly variable” populations from the T-REX comparison of day 7 to 0 for those with allergic asthma (n=6).

Supplemental table 3. Demographics and clinical data for subjects included in intracellular cytokine analysis.

| | **Cytokine profiling (n=6)** | **T cell phenotyping (n=32)** | **P-value[C]** |
|---|---|---|---|
| Age (years)[A] | 8.6 (6.3-11.5) | 6.0 (4.7-7.6) | 0.17[D] |
| Male sex, n (%) | 5 (83.3) | 23 (71.9) | >0.99[E] |
| White, n (%) | 3 (50.0) | 24 (75.0) | 0.64[E] |
| Total IgE (IU/ml)[A] | 78.2 (10.9-559.0) | 61.9 (34.1-112.3) | 0.822[D] |
| Atopic, n (%) | 4 (66.7) | 20 (62.5) | >0.99[E] |
| Medication, n (%) | | | |
| Inhaled corticosteroid (ICS) | 5 (83.3) | 24 (75.0) | >0.99[E] |
| Long-acting β-agonist | 5 (83.3) | 14 (43.8) | 0.08[E] |
| Antileukotriene | 2 (33.3) | 9 (28.1) | >0.99[E] |
| Oral prednisone | 0 (0.0) | 3 (9.4) | >0.99[E] |
| Dupilumab | 1 (16.7) | 2 (6.3) | 0.41[E] |
| Mepolizumab | 1 (16.7) | 1 (3.1) | 0.29[E] |
| Omalizumab | 0 (0.0) | 1 (3.1) | >0.99[E] |
| BAL phenotype, n (%) | | (n=31) | 0.75[E] |
| Isolated eosinophilia | 0 (0.0) | 2 (6.5) | |
| Isolated neutrophilia | 1 (16.7) | 9 (29.0) | |
| Mixed granulocytes | 1 (16.7) | 2 (6.5) | |
| Pauci-granulocytic | 4 (66.7) | 18 (58.1) | |
| Infection status, n (%) | | | |
| Rhinovirus | 0 (0.0) | 10 (31.3) | 0.17[E] |
| Other virus[B] | 0 (0.0) | 3 (9.4) | >0.99[E] |
| Any bacteria | 0 (0.0) | 10 (31.3) | 0.17[E] |
| Co-infection | 0 (0.0) | 6 (18.8) | 0.56[E] |

[A] Geometric mean (95% CI). [B] Adenovirus and Metapneumovirus. [C] Comparison of groups. [D] Mann-Whitney test. [E] Fisher's exact test.

Supplemental table 4. Spectral flow cytometry panel for intracellular cytokine analysis.

| Marker | Color | Clone | Vendor | Catalog number | Dilution |
|---|---|---|---|---|---|
| Viability | Live/Dead Blue | | Invitrogen | L23105 | 1:4000 |
| CD3 | BUV395 | SK7 | BD | 564001 | 1:50 |
| CD4 | BUV496 | OKT4 | BD | 750980 | 1:50 |
| CD45RO | BUV563 | UCHL1 | BD | 748369 | 1:100 |
| CXCR3 | BUV661 | 1C6/CXCR3 | BD | 741649 | 1:50 |
| CCR5 | BUV737 | 2D7/CCR5 | BD | 612808 | 1:25 |
| CD69 | BUV805 | FN50 | BD | 748763 | 1:100 |
| CD14 | BV510 | M5E2 | Biolegend | 301842 | 1:20 |
| CD19 | BV510 | HIB19 | Biolegend | 302242 | 1:20 |
| IL-4 | BV605 | MP4-25D2 | Biolegend | 500828 | 1:40 |
| CD103 | BV711 | Ber-ACT8 | Biolegend | 350221 | 1:50 |
| TNF-α | BV750 | MAb11 | BD | 566359 | 1:200 |
| IL-17A | BV785 | BL168 | Biolegend | 512338 | 1:100 |
| IL-13 | FITC | B-P6 | Invitrogen | BMS133FI | 1:50 |
| CD8 | PerCP | SK1 | Biolegend | 344708 | 1:200 |
| Granzyme B | RB705 | GB11 | BD | 570275 | 1:100 |
| IL-21 | PE | 3A3-N2.1 | BD | 562042 | 1:50 |
| CD161 | PE-Dazzle 594 | HP-3G10 | Biolegend | 339940 | 1:20 |
| IL-2 | PE-Cy7 | MQ1-17H12 | BD | 560707 | 1:50 |
| IL-5 | APC | JES1-39D10 | Miltenyi Biotec | 130-127-364 | 1:20 |
| IFN-γ | Alexa Fluor 700 | 4S.B3 | Biolegend | 502520 | 1:1000 |
| IL-22 | APC-Fire 750 | 2G12A41 | Biolegend | 366713 | 1:50 |

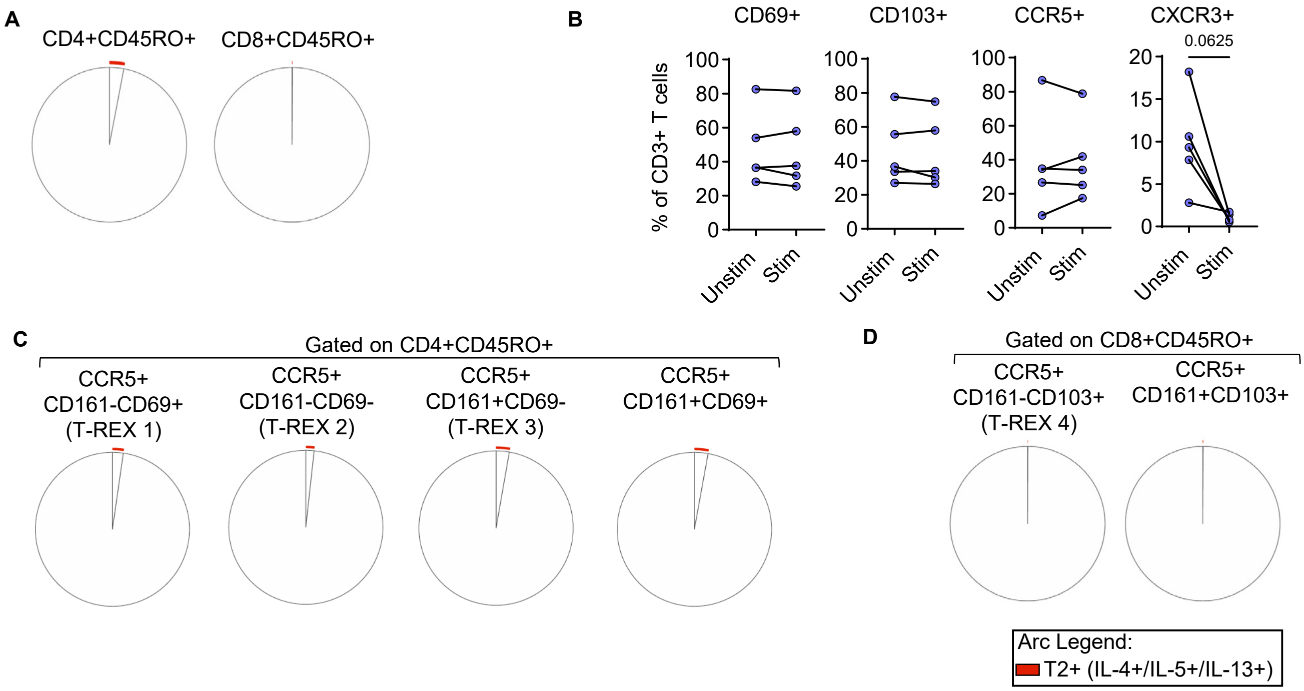


**Supplemental Figure 11. Type 2 cytokine production is minimal in the lower airways of children with recurrent wheeze.** (A) Spice plots showing the average frequency of cells positive for type 2 cytokine within memory CD4+ and CD8+ T cells (n=6). (B) Comparison of receptor-positive cell frequency in unstimulated and stimulated BAL cells as a percentage of CD3+ T cells (n=5). (C and D) SPICE plots showing the average frequency of cells positive for type 2 cytokines within each T cell population (n=6). Wilcoxon matched-pairs signed rank test used to calculate significance in A.

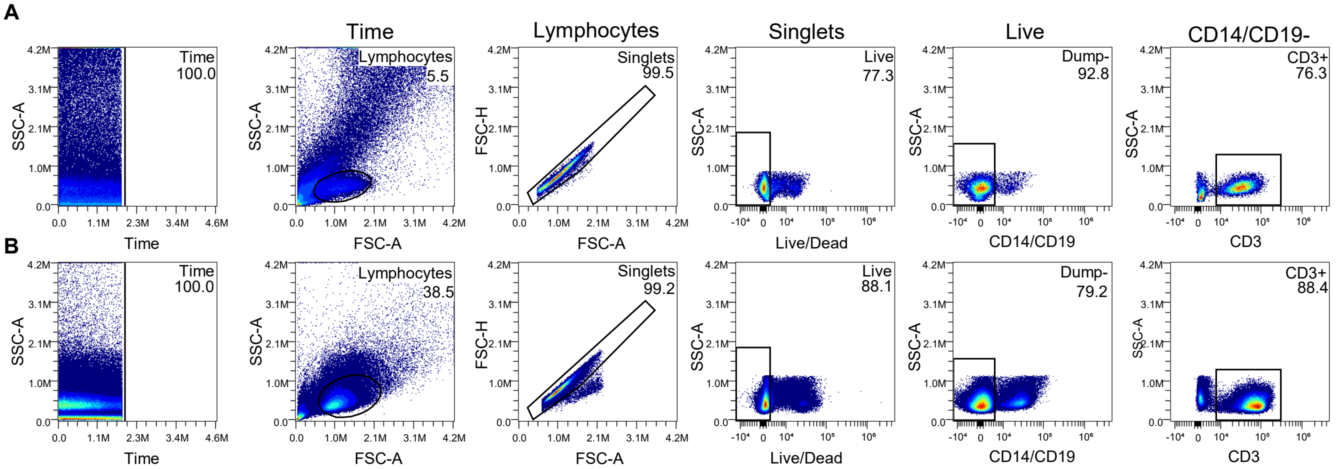


**Supplemental Figure 12. Representative scatter plots showing manual gating strategy for pre-processing of (A) blood and (B) BAL samples.**

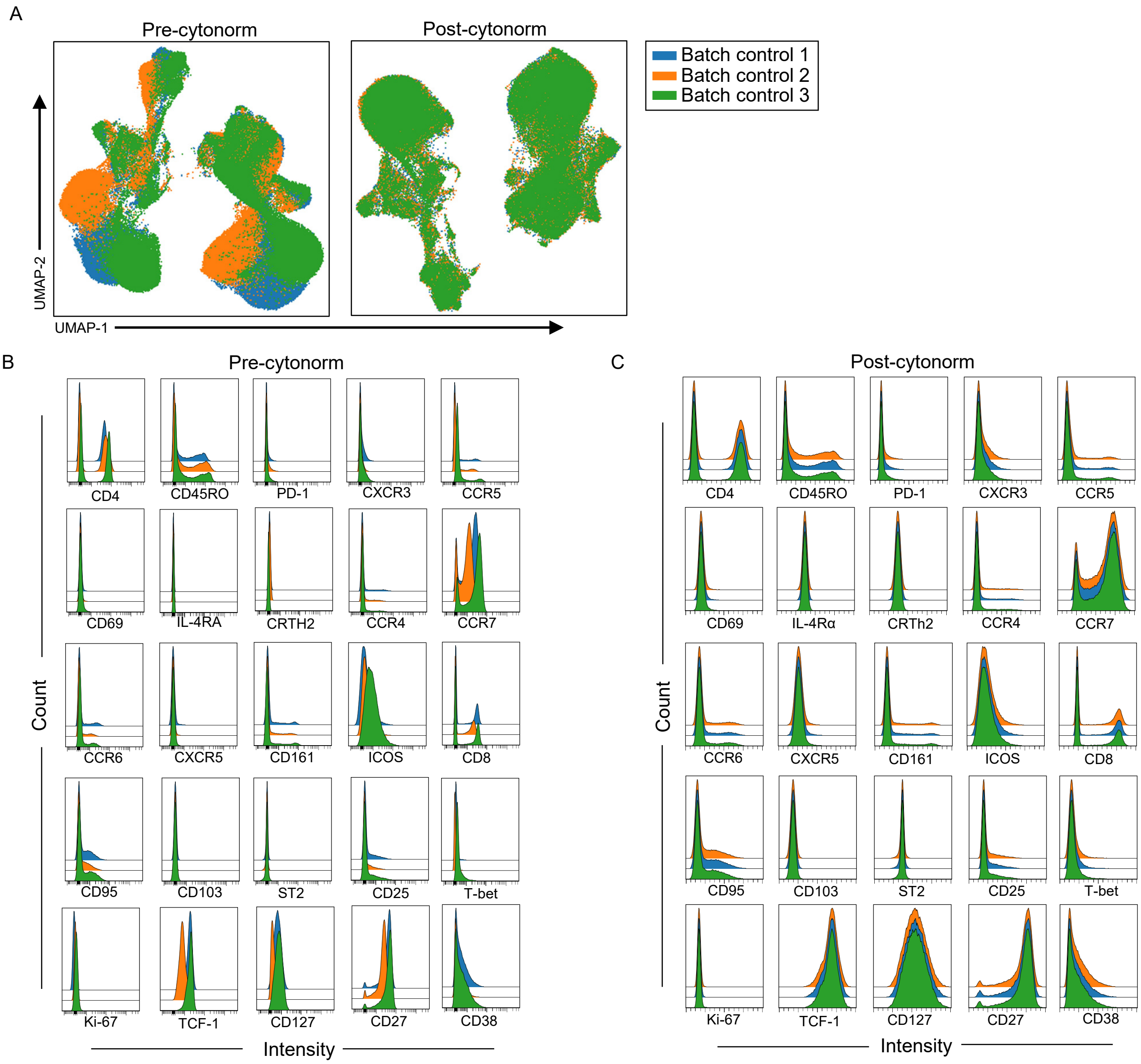


**Supplemental Figure 13. Results of normalization of blood and BAL CD3+ cells from 32 children with refractory wheeze.** (A) UMAP of batch control samples, pre- and post-normalization using Cytonorm. Histograms showing the distribution of each marker within each batch control, (B) pre- and (C) post-normalization.
